# The conformational distribution of a major facilitator superfamily peptide transporter is modulated by the membrane composition

**DOI:** 10.1101/2020.01.09.900316

**Authors:** Tanya Lasitza Male, Kim Bartels, Felix Wiggers, Gabriel Rosenblum, Jakub Jungwirth, Hagen Hofmann, Christian Löw

**Author notes:** Corresponding authors: Christian Löw, Hagen Hofmann. Tanya Lasitza Male and Kim Bartels contributed equally to this work.

## Abstract

While structural biology aims at explaining the biological function of membrane proteins with their structure, it is unclear how these proteins are modulated by the complex lipid composition of membranes. Here, we address this question by mapping the conformational distribution of the bacterial oligopeptide transporter DtpA using single-molecule fluorescence spectroscopy. We show that DtpA populates ensembles of conformers that respond sensitively to the environment. Detergents trap the transporter in an inward-open ensemble in which the substrate binding site faces the cytosol. However, reconstitution in Saposin nanoparticles with different lipid compositions, reveal a plethora of alternative conformations, including a fully inward-open ensemble whose existence had not been anticipated before. The relative abundance of these ensembles depends on the lipid composition of the nanoparticles. Our results therefore demonstrate that membranes sensitively affect the structural distribution of DtpA and we expect this to be a general property of membrane proteins.

## Introduction

Membrane proteins with their ability to shuttle, pump, exchange, and transmit signals across membranes^1-7^, represent nearly one third of the proteins in living organisms^8,9^. Lipid bilayers do not only compartmentalize cells, but the diverse functions of membrane proteins turn lipid bilayers into specialized and highly selective barriers for a broad variety of compounds. Lipids have been shown to play crucial roles in maintaining the structural and functional integrity of membrane proteins^10-15^. Unfortunately, biochemical and structural studies by x-ray crystallography or cryo-electron microscopy often require extracting these proteins from their natural environment^16-18^. For example, numerous structures of membrane proteins have been determined in detergent solution^19-22^. However, to derive mechanistic models of their function, it is essential to study these proteins in environments that mimic lipid bilayers as closely as possible^23-26^. Here we show that this is particularly important for a transporter belonging to the major facilitator superfamily (MFS).

The MFS is one of the largest families of membrane transporters in nature^27,28^. They mediate the uptake of a broad variety of substrates and play a crucial role in cell homeostasis^29^. Members of the MFS share a common fold^30^: they are comprised of twelve transmembrane helices that are organized in two six-helix bundles, the N-terminal and the C-terminal domain (Fig. 1, 2a). Residues from both domains form the substrate-binding site in the center of the protein.

**Fig 1.**
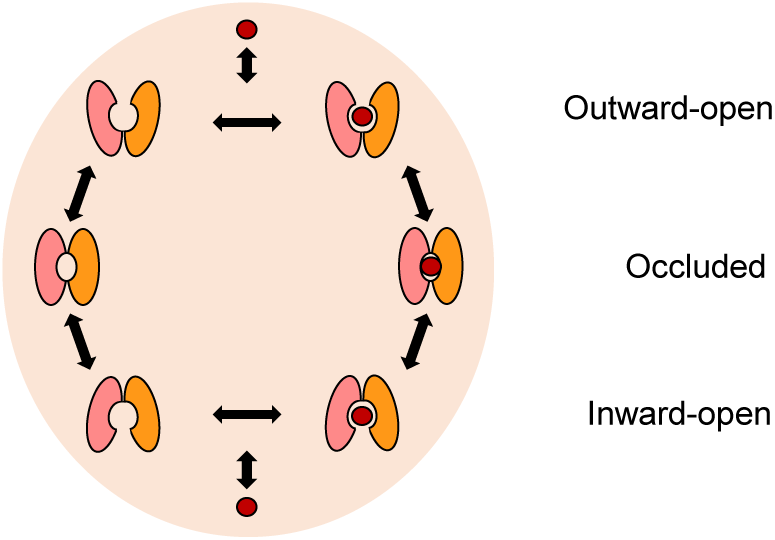
Conformational variety of the MFS transport cycle. The transporter binds the substrate in the outward-open conformation and shuttles it through the intermediate occluded state to the cytoplasmic side where it is released from the inward-open state.

**Fig 2.**
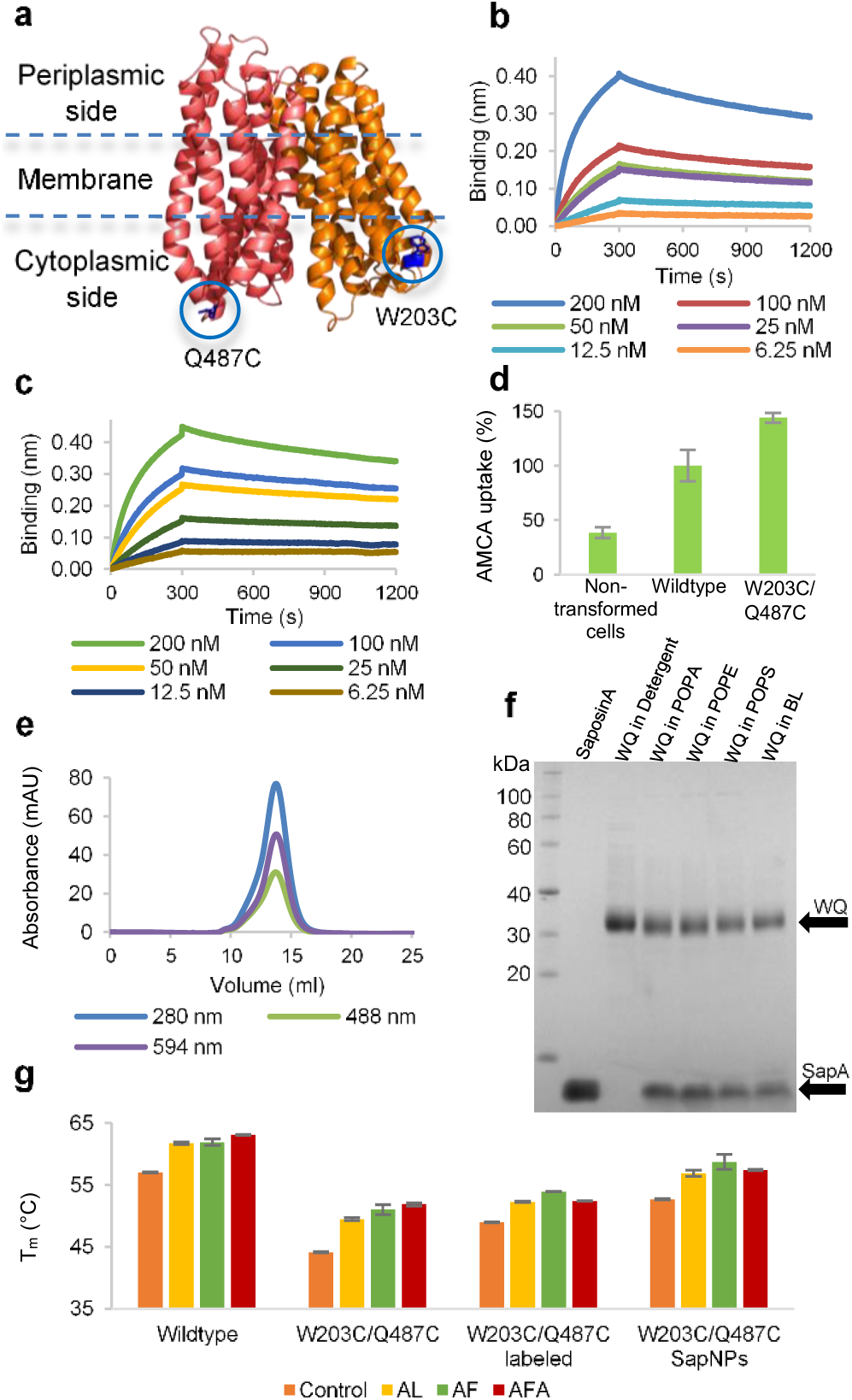
Confirming the structural and functional integrity of the DtpA variants used for smFRET. (a) Labeling positions of the DtpA variant WQ (W203C/Q487C) in the N-terminal (orange) and C- terminal domain (pink). (b) Bilayer interferometry, showing association and dissociation kinetics of nanobody binding to the WQ variant. Color code indicates the concentration of WQ. (c) Same as b for the FRET- labeled WQ variant. (d) *In vivo* uptake activities of DtpA wildtype and the WQ variant. Experiments were performed in triplicates. (e) Size exclusion (SEC) profile of the WQ variant after labeling, confirming the absence of oligomeric or aggregated species. (f) SDS-PAGE of WQ before and after reconstitution into SapNPs with various lipids confirming SapNPs assembly by co-elution of Saposin A and reconstituted WQ. (g) NanoDSF thermal stability profile of wildtype DtpA, labeled WQ and unlabeled WQ, and WQ incorporated in POPE SapNPs in the absence and presence of the ligands Ala-Leu (AL), Ala-Phe (AF) and Ala-Phe-Ala (AFA). Experiments were performed in triplicates.

It has been suggested that MFS transporters function via an alternate access mechanism^31,32^ where transport is mediated through conformational changes that allow the active center to face either side of the membrane. A full transport cycle requires at least three different conformations^33^: inward-open, occluded, and outward-open (Fig 1). A more recent model based on available crystal structures, named Clamp-and-Switch model^34^, proposes that substrate binding in the outward-open state causes the tips of two membrane helices to close, which results in a partially occluded conformation. This clamping step is followed by a switching motion in which the N-terminal and C-terminal domains move against each other, thus forming the inward-open state with the substrate-binding site being exposed to the cytoplasm.

A particularly important subfamily of the MFS are proton-dependent oligopeptide transporters (POTs)^27^. POTs utilize proton gradients^35^ to uptake di- and tripeptides but also peptide-mimetic drugs^36,37^. Their remarkable substrate promiscuity^38,39^ together with their highly abundant MFS fold^40^ makes them ideal models to understand the interplay between conformational changes and transport. Numerous bacterial POT structures have been determined to date^41-58^. We focus on the *E. coli* peptide transporter DtpA^59,60^ to explore the effect of different environments such as detergents and Saposin nanoparticles^61,62^ (SapNPs) with different lipid compositions on the conformation of the protein using single molecule FRET (smFRET)^63-68^. DtpA is an ideal model system since its substrate specificity is remarkably similar to that of its homologs PepT1 and PepT2 from human^36,69-75^. In contrast to its human homologs, DtpA can be expressed and purified from bacterial expression systems and remains stable in detergent solution allowing to develop a straightforward site-specific labeling protocol with FRET-fluorophores.

## Results

### Preparation of DtpA variants for FRET experiments

To identify optimal labeling positions for the thiol-maleimide coupling^76^ of our FRET fluorophores, we performed an extensive screening of multiple DtpA variants. First, we replaced three internal cysteines by serine residues to avoid any unspecific labeling that could impair the functionality of the transporter. Although these mutations destabilize DtpA (Fig. S1), we found that the structural and functional integrity was preserved in two variants, which carry the labels on the cytoplasmic side: W203C/Q487C (WQ) and W203C/T351C (WT) (Fig. 2a, 2e, S2-3), thus making them optimal for our FRET experiments. Both variants maintained their binding affinity to known substrates (Fig. 2g, S1) and to a conformation-specific nanobody (N00) (Fig. 2b, S4-5, Table S1) that binds the periplasmic side of DtpA^57^. In addition, these variants are fully functional as probed by an *in vivo* transport assay using the fluorescent substrate N-7-amino-4-methylcoumarin-3-acetic acid coupled to β-Ala-Lys (AK-AMCA)^73^ (Fig. 2d, S6). Importantly, we also confirmed that labeling with donor (Alexa Fluor 488) and acceptor (Alexa Fluor 594), does not affect the structural and functional integrity of the transporter. Similar dissociation constants of N00 binding were observed for the wildtype protein (K_D_= 7.13±0.03 nM), the WQ variant (K_D_=6.28±0.03 nM), and the FRET labeled WQ variant (K_D_=5.40±0.03 nM), (Table S1, Fig. 2b-c, S4-5). The thermal stability profiles of labeled and unlabeled variants were similar in the presence or absence of substrates (Fig. 2g, S7) and, when reconstituted in SapNPs, the variants show an increased stability, as expected from a previous study^61^, (Fig. S8). DtpA variants could be efficiently reconstituted in SapNPs of various lipid compositions, and formed homogeneous particles as monitored by SEC and SDS-PAGE (Fig 2f, S9). Since both variants (WQ and WT) show very similar results, we mainly focus on the WQ variant in the following. However, complementary data for the WT variant are shown in the supplementary information (Fig S1-3, S5-7, S10-11, S15, S22).

### Detergents as membrane mimicking environment

Maltoside based non-ionic detergents such as DDM (n-dodecyl-β-D-maltoside) and LMNG (lauryl-maltose-neopentyl-glycol) are commonly used for structural and functional studies of membrane transporters such as DtpA^18,77,78^. However, all current x-ray structures show POTs in an inward-open conformation and the lack of outward-open structures is currently hampering an atomic level understanding of the transport cycle. We conjectured that the crystallization bias towards the inward-open conformation could indicate that outward-open conformations are thermodynamically disfavored in LMNG micelles. We therefore started our smFRET experiments with freely diffusing DtpA at a LMNG concentration above the critical micelle concentration (0.002%). The FRET histograms of both variants show two populations (Fig. 3a-b, S10a-b). We find a dominant peak at intermediate FRET efficiencies with an average value of *E* = 0.50±0.03 (Fig. 3a), which is in good accord with the expected value of 0.54 based on the x-ray structure if the conformational freedom of the dye linkers is taken into account (Fig. S11). A second minor conformation (≈ 12%) is found at high FRET efficiencies of *E* = 0.89±0.02. Based on the alternate access model (Fig. 1), high FRET values are indeed expected for outward-open or occluded conformations of DtpA, in which the cytoplasmic labeling sites will be in a closed state. To identify the high-FRET population species, we added a conformation-specific nanobody (N00) that binds the transporter on the periplasmic side. If the high-FRET population represents the outward-open conformation, we expect the high-FRET population to disappear at saturating concentrations of N00. However, despite its nanomolar affinity for DtpA as demonstrated in independent biolayer interferometry experiments (Fig. 2c, Table S1), the high-FRET species does not vanish, indicating that this population rather presents conformations that are sampled independently of N00-binding to the opposite periplasmic side. These results do not allow an unambiguous assignment of the high-FRET population to an occluded or outward-open state and rather imply that the periplasmic and cytoplasmic sides of DtpA retain a certain degree of independence in LMNG.

**Fig 3.**
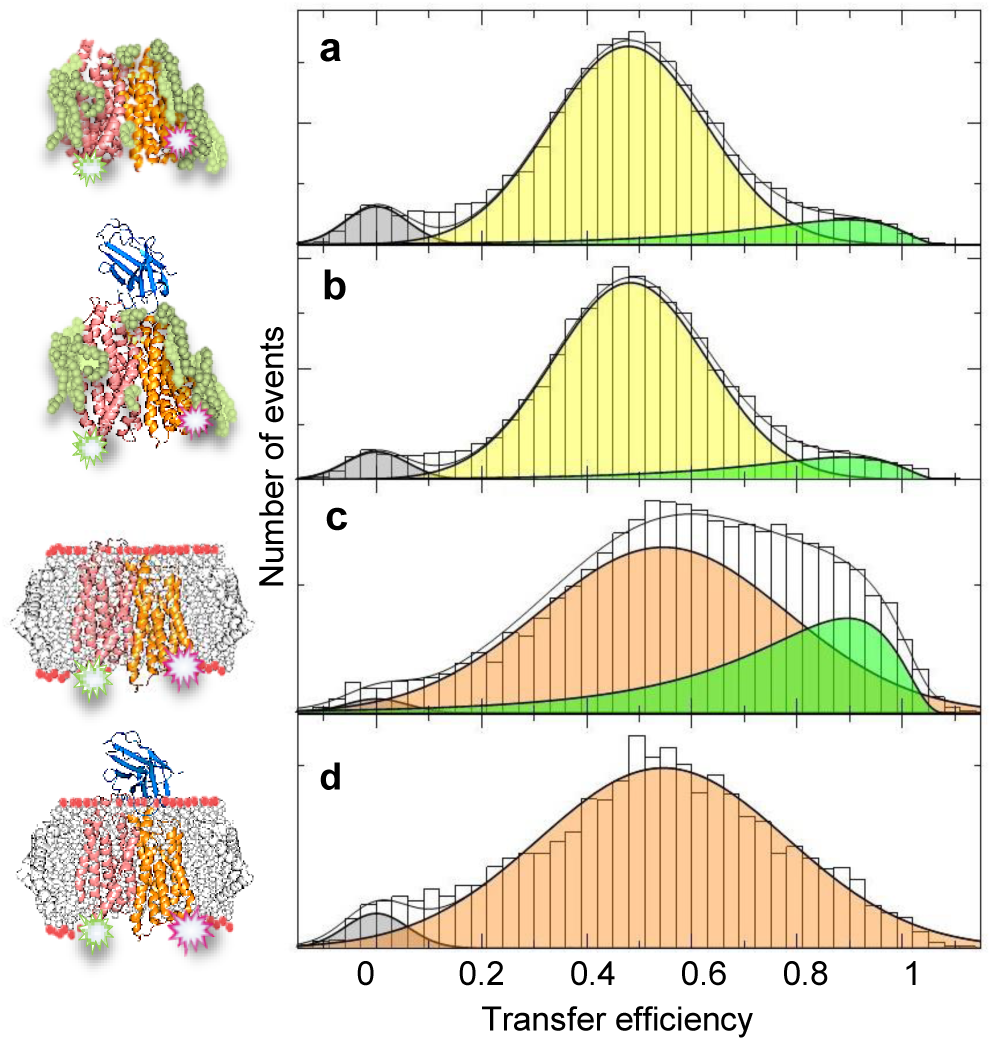
FRET efficiency histograms of DtpA variant WQ in LMNG detergent (a) and (b) and reconstituted in membrane mimicking POPE SapNPs (c) and (d). FRET histograms in (b) and (d) were obtained at saturating concentrations (8 µM) of nanobody (N00). Solid lines are fits with a superposition of Gaussian and log-normal peaks to identify the position of high-FRET (green) and intermediate FRET (yellow-LMNG, orange-SapNPs) peaks. The peak at zero transfer efficiency (gray) results from molecules without an active acceptor dye. The schematics (left panel) illustrates the different environments probed in the experiments with LMNG (green), nanobody (blue), SapNP-lipids (gray), and DtpA (color as in Fig. 2a). The labeling positions are indicated on the x-ray structure of DtpA.

In summary, we found that DtpA in LMNG micelles preferentially adopts an inward-open ensemble, which indeed explains the strong bias of current x-ray structures towards the inward-open structure. Any effector that shifts this equilibrium towards outward-open or occluded states provides a chance to obtain high-resolution x-ray structures of these missing states. We therefore checked the effect of ten different substrates on the balance between high-and low-FRET populations. Even though the substrates efficiently bind to the transporter (Fig. S12), there is no change in the observed FRET histograms (Fig. S13). Since DtpA is a proton-substrate symporter, we also recorded smFRET histograms at pH-values in the range between pH 4 to 8. However, the FRET distributions also remained unaltered by changes in pH (Fig. S14). We therefore conclude that the inward-open conformer of DtpA is very stable in LMNG and cannot easily be shifted to alternative conformations, a finding that is in accord with previous x-ray structures that only spotted marginal changes in DtpA in the prodrug bound and apo-state^57^.

### Conformations of DtpA in SapNPs

However, detergent micelles do not reflect the native environment of a cell membrane. To obtain information on the conformational ensemble of DtpA under more native conditions, we performed experiments in Saposin A nanoparticles (SapNPs) with different lipid compositions. We first reconstituted DtpA WQ in SapNPs using POPE (1-Palmitoyl-2-oleoyl-sn-glycero-3-phosphoethanol-amine), the most abundant phospholipid in *E. coli*^.79^ The FRET histogram of DtpA WQ in POPE SapNPs differs substantially from that obtained in LMNG micelles (Fig. 3c). The distribution is broader and shows an increased population at high FRET. Contrary to LMNG, the addition of N00 efficiently suppresses this high-FRET population (Fig. 3d). In line with the alternate access model, this result indicates a strong coupling between the cytoplasmic and periplasmic side of DtpA, suggesting that the high-FRET population in POPE SapNPs in the presence of N00 can be assigned to the outward-open state of DtpA. However, even at saturating concentrations of N00 (Fig. 3d), the FRET-distribution is significantly broader than that obtained in LMNG (Fig. 3b and 3d). Broadening of smFRET histograms can originate from a variety of sources including rotational freedom of the dyes, i.e., high fluorescence anisotropy^80^, static quenching due to contacts between FRET-dyes, and aromatic amino acids^81^, and conformational heterogeneity. Since we are particularly interested in the latter origin, we performed control experiments to rule out that the broadening results from anisotropy or quenching effects. We found steady-state donor fluorescence anisotropies of *r* = 0.219±0.156 in the absence and *r* = 0.219±0.154 in the presence of N00. Since similar values are found in LMNG micelles, the histogram broadening in SapNPs does not result from a restricted orientational freedom of the FRET-dyes (Fig. S15). In addition, using nanosecond fluorescence correlation spectroscopy^65^ (nsFCS), we find that static quenching of the acceptor dye, which has the strongest effects on FRET histograms^81^, has only a marginal amplitude (Fig. S16) and is therefore unlikely to cause the enormous broadening of the FRET distribution. The most likely cause of the broad FRET distribution is therefore structural heterogeneity at timescales slower than the molecule diffusion time through the confocal spot in our microscope (∼ 1 ms), indicating that the inward-open conformation of DtpA in POPE SapNPs is rather an ensemble of states with a distance distribution than a single state with a well-defined distance. This heterogeneity is expected to be static at the fast nanosecond time-scale of the donor fluorescence lifetime^65^, which is expected to cause deviations from the predicted linear dependence between donor lifetime and transfer efficiency^65,66^. This deviation is indeed observed experimentally (Fig S17). Surprisingly, the same deviation is also found for DtpA in LMNG (Fig. S17), indicating that structural heterogeneity is also found in LMNG micelles. Hence, the narrower FRET distribution in LMNG arises from a faster sampling of this distribution compared to POPE SapNPs. Apparently, the lipid environment of DtpA does not only affect the distribution of structural states but it also affects dynamics. In summary, our results show that (i) the native membrane environment restores the coupling between cytoplasmic and periplasmic side and (ii) it slows the dynamics within the inward-open ensemble.

### Phospholipid compositions tune the ensemble of DtpA conformers

After demonstrating that a native-like membrane environment causes significant changes in the conformation and dynamics of DtpA compared to LMNG micelles, we checked whether DtpA is also affected by the type of phospholipids. We therefore reconstituted labeled DtpA WQ in SapNPs composed of POPS (1-palmitoyl-2-oleoyl-sn-glycero-3-phospho-L-serine), POPA (1-palmitoyl-2-oleoyl-sn-glyc-ero-3-phospho-L-alanine), or BL (brain lipid extract) (Fig. 4 a-d). Indeed, the FRET histograms differ strongly in the four membrane environments. Up to three populations can be distinguished. The high- and intermediate FRET populations are identical to those found in POPE. However, an additional low-FRET peak (*E* = 0.25±0.02) is found in POPS and POPA (Fig. 4a-b), indicating a conformational ensemble with an even wider opening on the cytoplasmic side compared to the known x-ray structure of DtpA. In BL extracts on the contrary, neither the high-FRET nor the low-FRET peak is observed and the transporter samples exclusively the inward-open con-conformation (Fig. 4c). Similar to our results in POPE, the addition of N00 stabilizes the inward-open ensemble with intermediate FRET while all other conformations disappear (Fig. 4e-h), thus implying a structural coupling between the cytoplasmic and periplasmic side of DtpA. These results demonstrate that DtpA also exhibits a strong sensitivity to the specific lipid composition and it is currently unclear whether this sensitivity has also functional effects. The low-FRET population observed in POPS and POPA is particularly surprising. Its existence had not previously been anticipated by any transport model and its functional role in the transport cycle is currently unknown. We therefore checked whether the addition of substrate alters the abundance of the three populations. However, similar to our findings in LMNG, we found that the addition of substrates does not affect the conformational variety of DtpA (Fig. s18). Although the prodrug bound x-ray structure also does not show substantial structural changes compared to the apo-structure^57^, the invariance of the conformational distribution of DtpA to substrate in SapNPs is surprising. However, in cells, substrate concentrations differ significantly between periplasm and cytoplasm, thus generating an asymmetry that is not reproduced in our experiments. If the affinities for substrates are similar on both sides of DtpA, the isotropic concentration of substrate on both sides in our experiments may indeed cause the conformational invariance of DtpA towards substrate. To identify alternative parameters that can shift the conformational equilibrium we tested the thermal response of DtpA in LMNG and POPS SapNPs.

**Fig 4.**
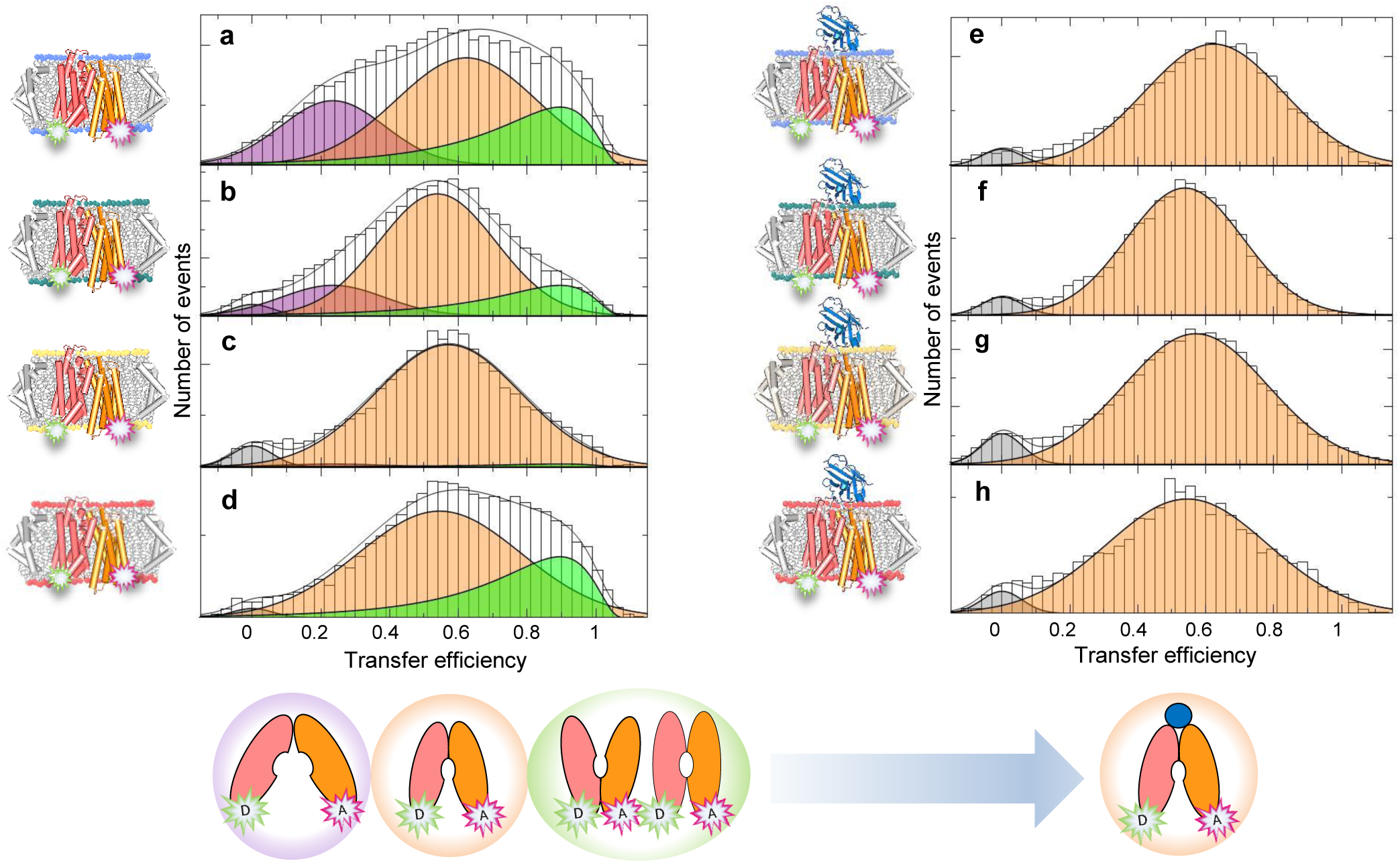
FRET efficiency histograms of the variant WQ in SapNPs of various phospholipid compositions in the absence (left, a-d) and presence of 8 µM concentrations of N00 (right, e-h). From top to bottom: POPS (a, e), POPA (b, f), BL (c, g), POPE (d, h). Solid lines are fits with a superposition of Gaussian and Log-normal functions. The relative populations with high FRET (green), inter-mediate FRET (orange) and the low FRET (purple) change upon addition of N00. The population of molecules without an active acceptor is shown in gray. Bottom: the mixture of conformational states in the absence of N00 collapses into the inward-open ensemble in the presence of N00.

### Temperature modulates the structural ensembles of DtpA in micelles and membranes

The experiments were performed in the temperature regime between 3°C and 30°C. We verified independently that the structural integrity of WQ in SapNPs and in LMNG is unaffected in this temperature range (Fig. 2g, S1, S7-8). In LMNG micelles, the inward-open population shifts continuously to lower transfer efficiencies with decreasing temperature (Fig. 5, left). This shift is also associated with a significant broadening. Clearly, more open-conformers are sampled at lower temperatures and the continuous transition confirms our interpretation that the cytoplasmic side of DtpA populates a broad ensemble of conformers. The broadening of the distribution hints at a slowing-down of the dynamics with which DtpA samples these conformations, which is indeed expected with decreasing temperatures. Importantly, the temperature-induced shift to lower FRET-values is even observed in the presence of saturating amounts of N00 in LMNG micelles (Fig. 6), thus confirming that the cytoplasmic side retains flexibility despite the bound N00 on the periplasmic side. Interestingly, the fraction of the unidentified high-FRET population is only marginally altered by temperature, which may indicate that it represents an extreme conformational tail of the inward-open ensemble rather than a defined state that is separated by a free energy barrier from the inward-open distribution. Importantly, a strong temperature-sensitivity is also observed in POPS SapNPs. Fits with partially constraint Gaussian and log-normal functions^82^ indicate a stabilization of the high-FRET population, i.e., the outward-open ensemble (Fig. 5 right). In contrast, the balance between the two low-FRET populations remains unaffected by temperature (Fig. 5). Although these interpretations depend on the precise fitting model - we reinforced identical positions of the high- and low-FRET population for all temperatures - our experiments show a clear shift towards higher transfer efficiency with increasing temperature in both LMNG micelles and POPS SapNPs. At the physiological habitat temperature of *E. coli* (37°C), this trend is likely to populate the outward-open (high-FRET) species to a significant amount.

**Fig 5.**
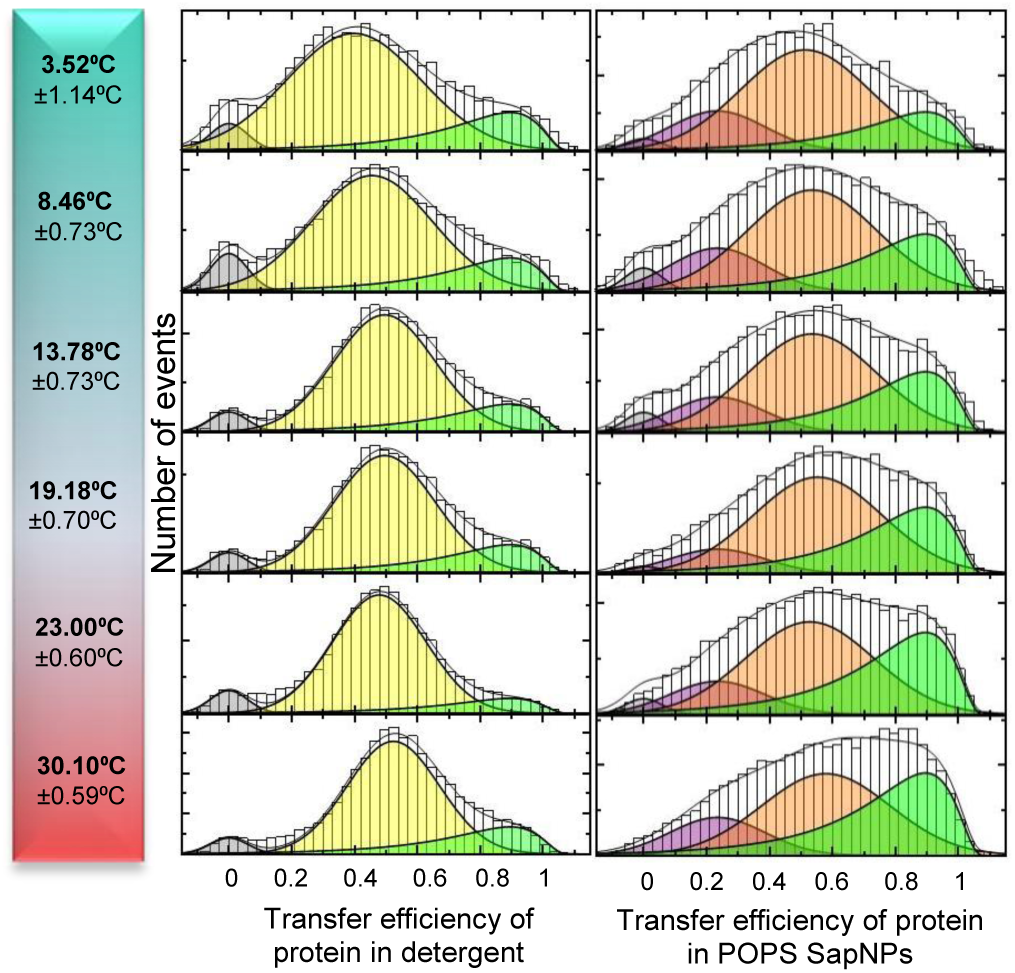
FRET histograms of DtpA (WQ) in LMNG (left) and membrane mimicking POPS SapNPs (right), as a function of temperature (indicated). The color code is identical to Fig. 2 and 3.

**Fig 6.**
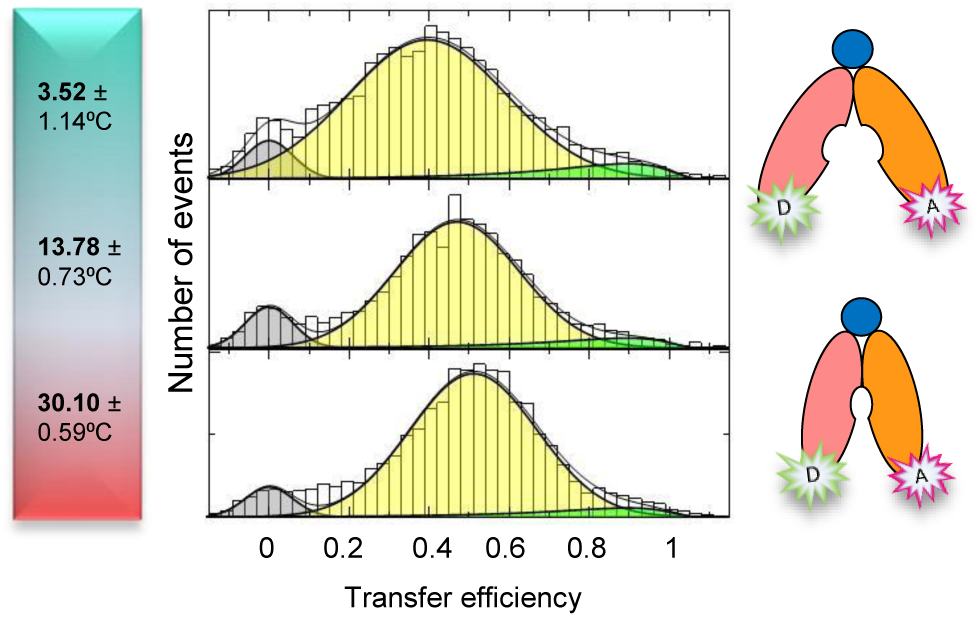
Effect of temperature on the FRET histograms of DtpA (WQ) in LMNG and in the presence of N00 (8 µM) (left). (Right) Scheme to illustrate the conformational independence between cytoplasmic and periplasmic side.

## Discussion

POTs have been extensively studied by x-ray crystallography and biochemical transport assays over the past years^27,36,39,59,70,73,83,84^. Numerous structures of various bacterial homologues in the absence and presence of substrates, drugs, and prodrugs are available, highlighting crucial residues for substrate binding and proton coupling^41-58^. Unfortunately, only inward-open and partially inward-open states of POTs have so far been described at atomic resolution and the influence of the membrane environment on structure and transport has been neglected^41,42,47,52,53,55,58^. Our aim was to close this gap by mapping the conformational distribution of POTs in different membrane-mimicking environments using smFRET. Our results show that the conformational distribution of the bacterial POT DtpA depends sensitively on two major parameters: (i) lipid environment and (ii) temperature. In particular, we show that both conformational distribution and dynamics differ substantially between micelle and a lipid membrane environment. Hence, care has to be taken when interpreting functional aspects based on x-ray structures obtained in detergent. Clearly, the precise lipid composition in *E. coli* is difficult to reproduce *in vitro* and at best, our experiments covered extreme limits. However, given the complexity of natural membranes in which compositions can vary on sub-micrometer length scales, e.g., in micro-domains^85^, the heterogeneity found among four membrane-like environments in our experiments may suggest that DtpA is also conformationally heterogeneous in the cell membrane. While x-ray structures picture models with well-defined states, our results rather indicate that such states are better described as structural ensembles that exhibit significant dynamics. The role of these dynamics on functionally relevant timescales in a productive transport cycle has to be the topic of future experiments.

## Methods

### Chemicals

Unless specified otherwise, chemicals were purchased from Sigma-Aldrich, phospholipids and membrane extracts from Avanti Polar Lipids, Inc., n-dodecyl-β-D-maltoside (DDM) and lauryl-malt-ose-neopentyl-glycol (LMNG) from Anatrace, antibiotics, iso-propyl-β-d-thiogalactopyranoside (IPTG) and 1,4-dithiothreitol (DTT) from Roth, DNase I from Appli-Chem, Lysozyme and protease inhibitor cocktail from Roche, di/tripeptides were purchased from Sigma-Aldrich, Fluka, and Bachem, Alexa Fluor 488 C5 maleimide and Alexa Fluor 594 C5 maleimide from ThermoFisher and Fluka and Terrific broth (TB) from Melford.

### Protein Constructs

DtpA variants (Uniprot ID: P77304) were cloned into the pNIC-CTHF vector (Addgene plasmid-ID Plasmid #26105) using LIC cloning^86^. Point-mutations for FRET labeling variants were generated by site-directed mutagenesis. Briefly, wildtype DtpA in pNIC-CTHF vector was used as template for PCR with mutagen primers containing the desired mutation (Table S2). Mutations were confirmed by DNA sequencing. Saposin A was c cloned into pNIC-Bsa4 expression vector Addgene plasmid-ID Plasmid #26103) using LIC cloning. The nanobody (N00) was cloned into a pMESy2 vector.

### DtpA Variants, Expression Purification and Labeling

DtpA variants were expressed and purified as previously described^57^. The constructs were transformed into *E. coli* strain C41(DE3) and grown in TB medium supplemented with 30 µg/ml kanamycin at 37°C. The cells were induced by 0.2 mM IPTG at OD_600 nm_ of 0.6, incubated further for 16 hours at 18°C, and harvested by centrifugation (9379 g, 15 min, 4°C). Then the cells were resuspended in lysis buffer (20 mM sodium phosphate pH 7.5, 300 mM NaCl, 5% (v/v) glycerol, 15 mM imidazole, 0.5 mM TCEP, 5 units/ml of Dnase I, 1 mg/ml lysozyme, and protease inhibitor). Typically, 5 ml of lysis buffer were used for 1 g of cell pellet. Cells were lysed by three cycles using an EmulsiFlex-C3 (Aventin) at 10,000 psi. After a low-speed centrifugation step (10,000 g, 15 min, 4°C) that separated the undisrupted cells and debris, an ultracentrifugation step of the supernatant was performed to pellet the membrane fraction (142,400 g, 50 min, 4°C). The crude membranes were resuspended in lysis buffer, supplemented with 1% LMNG, 0.5 mM TCEP and protease inhibitor, and stirred for 60 min at 4°C. After an additional ultracentrifugation step (104,600 g, 50 min, 4°C), DtpA was purified by Immobilized Metal Affinity Chromatography (IMAC) using Ni-NTA agarose (ThermoFisher). His-tagged proteins were bound to the resin for 60 min at 4°C on a rotating wheel, extensively washed with increasing imidazole concentration (20 mM sodium phosphate pH 7.5, 300 mM NaCl, 5% (v/v) glycerol, 15-25 mM imidazole, 0.01% (w/v) LMNG, 0.5 mM TCEP) and eluted in elution buffer (20 mM sodium phosphate pH 7.5, 150 mM NaCl, 5% (v/v) glycerol, 250 mM imidazole, 0.01% (w/v) LMNG, 0.5 mM TCEP). The eluate was concentrated to 1 ml using a 50 MWCO concentrator (Corning Spin-X UF concentrators) and incubated with 10 mM DTT for 30 min at 4°C. Size exclusion chromatography (SEC) was performed on an ÄKTA Pure system (GE Healthcare Life Sciences) using a Superdex 200 Increase 10/300 GL column (GE Healthcare Life Sciences) equilibrated with SEC Buffer (20 mM sodium phosphate pH 7.5, 150 mM NaCl, 5% (v/v) glycerol, 0.01% (w/v) LMNG). Fractions containing the protein were pooled and concentrated to 500 µl. For labeling, the variants were incubated for 2 hours at RT under gentle agitation with a 1:1 mix of Alexa Fluor 488 C5 maleimide and Alexa Fluor 594 C5 maleimide dyes (ThermoFisher) in a 1:2.4 molar ratio of protein to dyes. To stop the labeling reaction, the samples were incubated with 1 mM L-Glutathione for 30 min at RT. Free dyes were removed using a PD-10 desalting column (GE Healthcare Life Sciences). An additional IMAC was performed, the eluate from the desalting column was incubated with Ni-NTA agarose for 60 min at 4°C on a rotating wheel, extensively washed with increasing imidazole concentration, and eluted in elution buffer. TEV protease was added to the eluate 0.3 mg for 0.5 l culture. The sample was dialyzed overnight at 4°C against SEC buffer. Negative IMAC was performed to recover the cleaved labeled variants. Protein containing fractions were loaded on SEC Superdex 200 Increase 10/300 GL column (GE Healthcare Life Sciences) equilibrated with SEC buffer (20 mM sodium phosphate pH 7.5, 150 mM NaCl, 5% (v/v) glycerol, 0.01% (w/v) LMNG). Fractions containing the protein were pooled and concentrated.

### Nanobody (N00) Expression and Purification

The nanobody selection was previously described^57^. The nanobody expression plasmid was transformed into the *E. coli* strain WK6. The cells were grown at 37°C in TB medium supplemented with 100 µg/mL Carbenicillin. At an OD_600 nm_ of 0.7 the cells were induced with 1 mM IPTG and incubated for 16 h at 27°C. Cells were harvested (9379 g, 15 min, 4°C). Afterwards, the cells were resuspended in 5 ml TES buffer (200 mM Tris pH 8.0, 0.5 mM EDTA, 500 mM sucrose) per 1 g of pellet. Fourfold diluted TES buffer was added to perform an osmotic shock. Cell debris were removed by centrifugation (10,000 g, 15 min, 4°C) and the supernatant was recovered and applied on a Capture Select column (ThermoFisher), equilibrated previously with wash buffer (20 mM sodium phosphate pH 7.5, 150 mM NaCl), and the protein was eluted in an elution buffer (20 mM sodium phosphate pH 7.5, 150 mM NaCl, 2 M MgCl2). The eluted fractions were loaded on SEC HiLoad 16/600 Superdex 75 pg column (GE Healthcare Life Sciences), equilibrated with Nanobody Wash Buffer. Fractions containing protein were pooled and concentrated using a 5 MWCO concentrator (Corning Spin-X UF concentrators).

### Saposin A (SapA) Expression and Purification

The SapA expression plasmid was transformed into the *E. coli* strain Rosetta gami-2(DE3). Transformed bacteria were grown at 37°C in TB medium supplemented with 30 µg/mL kanamycin, 34 µg/mL chloramphenicol and 10 µg/mL tetracycline. At an OD_600 nm_ of 0.7, the cells were induced with 1 mM IPTG and incubated for 4 h at 37°C. Cells were harvested by centrifugation (9379 g, 15 min, 4°C). Typically, 1 g of pellet was suspended in 5 ml lysis buffer (20 mM sodium phosphate pH 7.5, 300 mM NaCl, 5% (v/v) glycerol, 15 mM imidazole, supplemented with 5 units/ml DnaseI, 1 mg/ml lysozyme and protease inhibitor). Cells were lysed by three cycles using an EmulsiFlex-C3 (Aventin) at 10.000 psi. The lysate was incubated at 80°C, 10 min. Non-lysed cells, debris, and aggregated proteins were removed by centrifugation (43,670 g, 15 min, 4°C). The supernatant was purified by IMAC using Ni-NTA agarose. His-tagged samples were bound to the resin for 60 min at 4°C on a rotating wheel, extensively washed (20 mM sodium phosphate pH 7.5, 300 mM NaCl, 5% (v/v) glycerol, 15-30 mM imidazole), and eluted in an elution buffer (20 mM sodium phosphate pH 7.5, 150 mM NaCl, 5% (v/v) glycerol, 400 mM imidazole). TEV protease was added to the eluate and 2 mg were used to cleave protein from 3 l culture. The sample was dialyzed overnight at 4°C against Saposin SEC Buffer (20 mM sodium phosphate pH 7.5, 150 mM NaCl). Negative IMAC was performed to recover cleaved protein. The last purification step was SEC on a HiLoad 16/600 Superdex 75 pg column equilibrated with Saposin SEC Buffer. Fraction containing protein were pooled, and concentrated using a 5 MWCO concentrator (Corning Spin-X UF concentrators).

### Cysteine Accessibility Assay

We tested the accessibility of cysteine residues of DtpA variants before and after labeling. Briefly, 0.15 mg/mL of each variant were incubated with methoxypolyethylene glycol maleimide 5000 (PEG-maleimide) (Sigma-Aldrich) at 2 mM final PEG-maleimide concentration for 30 min at RT. SDS-PAGE was performed using a 4-12% Bis-Tris gel (Expedeon) and stained using InstantBlue Coomassie Protein Stain (Expedeon).

### *In vivo* Uptake Assay

The uptake assay of the fluorescent N-7-amino-4-methylcoumarin-3-acetic acid coupled to β-Ala-Lys (AK-AMCA) dipeptide was performed as described previously with minor changes^73^. For *in vivo* uptake, the DtpA expression plasmid was transformed into *E. coli* strain C41(DE3). Transformed bacteria were grown at 37°C in TB medium supplemented with 30 µg/mL kanamycin. At an OD_600 nm_ of 0.6, cells were induced with 0.2 mM IPTG and incubated for further 3 h at 37°C. Cells were harvested by centrifugation at 3214 g for 15 min at 4°C and suspended in assay buffer (20 mM sodium phosphate pH 7.5, 150 mM NaCl, 5 mM glucose) to OD_600 nm_ of 10. In a final volume of 100 µl, 50 µl assay buffer, and 40 µl cells at OD_600 nm_ of 10 were incubated with 10 µl of 1 mM AK-AMCA for 20 min at 37°C. For the negative control, double distilled water instead of AK-AMCA was added. The reaction was stopped by the addition of 200 µl ice-cold assay buffer. Cells were washed twice with 200 µl ice-cold assay buffer and finally suspended in 200 µl ice-cold assay Buffer. Remaining fluorescence was measured using a Tecan Spark multimode microplate reader (Tecan Life Sciences) with an excitation of 350 nm and emission of 450 nm. To correct for the number of cells contributing to the fluorescence signal, the OD_600 nm_ for each sample was measured. Western blot was performed to account for differences in expression level of the variants and the wildtype. In short, SDS-PAGE using a 4-12% Bis-Tris gel (Expedeon) was performed and transferred on a PVDF membrane (Biorad). 2% BSA in TBST (Sigma) was used for blocking, TBST was used as washing buffer. The membrane was incubated with HisProbe-HRP conjugates antibody (ThermoFisher) for 1 h at RT. The blot was developed using Super Signal West Pico Substrate (ThermoFisher) and Super Signal West Femto Substrate (ThermoFisher) in a 1:10 ratio.

### Reconstitution of DtpA Variants in Saposin A Nanoparticles (SapNPs)

Labeled DtpA variants were reconstituted into SapNPs using a 1:20:35 molar ratio of DtpA variant:SapA:lipids^26^. Lipid stocks were prepared as previously described^26^. We used: 1-palmitoyl-2-oleoyl-sn-glycero-3-phospho-L-serine (POPS), 1-palmitoyl-2-oleoyl-sn-glycero-3-phosphoethanol-amine (POPE), 1-palmitoyl-2-oleoyl-sn-glycero-3-phosphate (POPA) and Brain Lipids Extract (Avanti Polar Lipids). Lipids were incubated at 37°C for 10 min, DtpA variant was added and incubated at RT for 15 min. After addition of SapA, the sample was incubated at RT for 20 min. Biobeads (Biorad) were added and the sample was incubated overnight at 4°C on a rotating wheel. The reconstituted DtpA was recovered by SEC Superdex 200 Increase 10/300 GL column (GE Healthcare), equilibrated with SEC buffer (20 mM sodium phosphate pH 7.5, 150 mM NaCl, 5% (v/v) glycerol).

### Thermal Shift Assay

The stability of DtpA was monitored by nanoDSF. 0.5 mg/mL of DtpA and its variants were incubated with the transporter substrates^73^: L-alanine-L-Leucine (AL), L-alanine-L-phenylalanine (AF), L-alanine-L-phenylalanine (AF) and L-alanine-L-phenylalanine-L-alanine (AFA) at 2.5 mM final ligand concentration for 10 min at RT. In addition, DtpA at 0.5 mg/ml was incubated with N00 at a molar ratio of 1:1.2 for 10 min at RT. Standard grade nanoDSF capillaries (Nanotemper) were loaded into a Prometheus NT.48 device (Nanotemper) controlled by PR. ThermControl (version 2.1.2). Excitation power was adjusted to 20% and samples were heated from 20°C to 90°C with a slope of 1 °C/min. All samples were examined in triplicates and and error bars represent standard deviations.

### Biolayer Interferometry

Dissociation constants (*K*_*D*_) of N00 binding to DtpA were measured using an Octet RED96 System (fortéBIO). To this end, N00 was biotinylated using an EZ-Link™ NHS-PEG4-Biotin kit (ThermoFisher). N00 was diluted to 5 μg/ml with octet buffer (20 mM sodium phosphate pH 7.5, 150 mM NaCl, 0.01% (w/v) LMNG) and loaded onto streptavidin (SA) biosensors (fortéBIO) that was hydrated with the same buffer. Unbound N00 was washed off with octet buffer. DtpA variants at 200 μM were then bound to N00. Experiments were performed at 25°C under shaking at 1000 rpm. Data were analyzed using Data Analysis v.10.0.3.1. software (fortéBIO) assuming a 1:1 stoichiometry of the DtpA-N00 complex. A Savitzky-Golay filter was applied to smooth the data.

### Rotational Isomeric State Model for FRET Efficiency Prediction

FRET prediction was done by first simulating possible dye positions 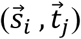 along with their probabilities 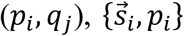 and 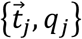, for donor and acceptor fluorophores. The FRET efficiency was then evaluated as 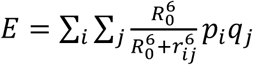, where 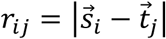 and *R*_0_ = 5.4 nm is the Förster radius. 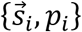 and 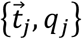 were obtained by modeling the fluorophores as bulky spheres, which were attached to the given C_β_ through a polyethylene chain of eleven monomers. The conformational freedom of the linker was accounted for using the rotational isomeric state model (RIS)^87^: each bond along the chain samples the anti, gauche+ and gauche-rotamers. Chains that sterically clashed^88^ with the protein were discarded. For computational ease, the conformational space was sampled stochastically as a biased random walk and probabilities for each step were derived from the RIS. Convergence was reached after sampling 10,000 linker conformations (standard deviation in predicted FRET <0.003). Geometric and energetic parameters of the linker were taken from Rehan, M *et al.*^87^, the fluorophore size from Kalinin, S. *et al.*^89^, and the DtpA structure from PDB entry 6GS7.

### Single Molecule FRET Microscopy

Single molecule FRET experiments were performed with a Micro-Time 200 confocal system (PicoQuant, Germany) equipped with an Olympus IX73 inverted microscope and two pulsed excitation sources (40 MHz) controlled by a PDL 828-L “Sepia II” (PicoQuant, Germany). Pulsed Interleaved Excitation (PIE)^90-92^ was used to identify molecules with active acceptor and donor dyes. Two light sources, a 485 nm Diode Laser (LDH-D-C-485, PicoQuant) and a white-light laser (Solea, PicoQuant) set to an excitation wavelength of 595 m were used to excite the donor and the acceptor dyes alternatingly. The laser intensities were adjusted to 100 μW at 485 nm and 20 μW at 595 nm (Pm100D, Thor Labs). The excitation beams were guided through a major dichroic mirror (ZT 470-491/594 rpc, Chroma) to a 60×1.2 NA water objective (Olympus) where they were focused into the sample. The samples were placed on quartz and glass coverslips (Esco Optics) with a 6 mm diameter cloning cylinder (borosilicate glass, Hilgenberg) glued on top. The experiments were conducted in fresh buffers containing 20 mM sodium phosphate, pH 7.5, 150 mM NaCl, 0.002 % LMNG and 20 mM DTT. The experiments with reconstituted variants in SapNPs were performed in the same buffer without the LMNG component. We verified that the addition of DTT does not significantly affect the structural integrity of SapNPs (Fig S23). The samples were diluted to 40-100 pM. Photons emitted from the sample were collected by the objective specified above. After passing the dichroic mirror (specified above), the residual excitation light was removed by a long-pass filter (BLP01-488R, Semrock) and the fluorescence light was focused on a 100 µm pinhole to remove out-of-focus light. Afterwards, the fluorescence photons were separated into donor and acceptor components using a dichroic mirror (585DCXR, Chroma, Rockingham, VT) and directed into donor and acceptor photon counting channels. Each light component was then focused onto the active area of a single-photon avalanche diode (SPAD) (Excelitas) after passing additional band-pass filters: FF03-525/50, (Semrock) for the donor and FF02-650/100 (Semrock) for the acceptor. For fluorescence anisotropy measurements, the emission light was first separated into its parallel and perpendicular components with respect to the linearly polarized excitation light via a polarizing beam splitter and each component was separated by two dichroic mirrors into donor and acceptor photons. The arrival time of the detected photons was recorded with a HydraHarp 400 time-correlated single photon counting (TCSPC) module (PicoQuant) and temperature controlled smFRET experiments were executed in a custom made system (Fig. S24), adapted from Nettels *et al.* and Aznauryan *et al.* ^93-95^. The temperature inside the confocal spot was determined by precisely measuring the diffusion coefficient of Oregon Green (ThermoFisher Scientific) in water using 2f-FCS^96^ (Fig. S25). The average of the fits from three independent measurements at each temperature were used to determine the viscosity of water given the hydrodynamic radius of Oregon Green, (0.6 nm)^96,97^. The known temperature dependence of the water viscosity^98^ was then used to calculate the temperature in the confocal spot. Errors in temperature (Fig. 5-6, S20-22) are standard deviations of the triplicates.

### SmFRET Data Analysis

As described previously^99,100^, photons from individual molecules, separated by less than 100 µs were combined into bursts if the total number of photons exceeded 80. Photon counts were corrected for background, acceptor direct excitation, and different detection efficiencies of the individual detectors^101^. To remove potential aggregates, we only included bursts with a burst duration < 3 ms and a PIE stoichiometry ratio *S* < 0.7^102^. FRET histograms were fitted with a combination of Gaussian and log-normal functions as previously described^82^.

### Fluorescence Lifetime Analysis

The average lifetimes of the donor in the presence of acceptor *τ*_*DA*_ (Fig. S17) is estimated by the mean arrival time of the photons relative to the exciting laser pulse. The expected value of *τ*_*DA*_ for a static inter-dye distance is given by: *τ*_*DA*_ = *τ*_*D*_(1 – *E*(*r*)), where *τ*_*D*_ = 4 ns is the lifetime of the donor in the absence of acceptor. In contrast, a distribution of distances *P*(*r*) alters this dependence^65^. As an upper limit for a broad distance distribution, we chose the Gaussian chain model^103^.

## Data availability

All data is available from the corresponding author upon reasonable request.

## Code availability

The code that was used to analyze the data is available from the corresponding authors upon request.

## Acknowledgments

We thank the Sample Preparation and Characterization facility at EMBL (Hamburg, Germany) for their support with nanoDSF and the biolayer interferometry measurements. We thank Dr. Haydyn Mertens EMBL (Hamburg, Germany) for generating the model of DtpA in SapNPs. We thank Prof. Phillip Selenko Weizmann Institute of Science (Rehovot, Israel), Prof. Jordan Chill, and Dr. Inbal Sher, Bar Ilan University (Israel) for the fruitful discussions concerning the reconstitution of membrane proteins in nanodiscs. Dr. Hofmann thanks the Benoziyo Fund for the Advancement of Science, the Carolito Foundation, the Gurwin Family Fund for Scientific Research, and the Leir Charitable Foundation. This work was funded by a joint grant from the German-Israeli Foundation (GIF grant #G-1288-207.9/2015) to Dr. Hagen Hofmann and to Dr. Christian Löw.

## Author Contributions

C.L. and H.H. designed the research and wrote the paper. T.L.M. and K.B. contributed equally to this work by designing research, performing experiments, analyzing data, and writing the paper. F.W. built the temperature controlled sample device for smFRET experiments. G.R. provided biochemical support. J.J. performed the rotational isomer model simulations.

## Additional information

Supplementary information is available for this paper.

## References

1 Robertis, D. Cell and molecular biology. 8th Edition. (Lea and Febiger, Philadelphia, PA; None, 1987).

2 Scherrmann, J. M. in Comprehensive Medicinal Chemistry II (eds John B. Taylor & David J. Triggle) 51–85 (Elsevier, 2007).

3 Ubarretxena-Belandia, I. & Engelman, D. M. Helical membrane proteins: diversity of functions in the context of simple architecture. Current Opinion in Structural Biology 11, 370–376, doi:https://doi.org/10.1016/S0959-440X(00)00217-7 (2001).

4 Rice, A. J., Park, A. & Pinkett, H. W. Diversity in ABC transporters: Type I, II and III importers. Critical Reviews in Biochemistry and Molecular Biology 49, 426–437, doi:10.3109/10409238.2014.953626 (2014).

5 Borges-Walmsley, M. I., Mckeegan, K. S. & Walmsley, A. R. Structure and function of efflux pumps that confer resistance to drugs. Biochemical Journal 376, 313–338, doi:10.1042/bj20020957 (2003).

6 Borges-Walmsley, M. I. & Walmsley, A. R. The structure and function of drug pumps. Trends in Microbiology 9, 71–79, doi:https://doi.org/10.1016/S0966-842X(00)01920-X (2001).

7 Hilger, D., Masureel, M. & Kobilka, B. K. Structure and dynamics of GPCR signaling complexes. Nature Structural & Molecular Biology 25, 4–12, doi:10.1038/s41594-017-0011-7 (2018).

8 Wallin, E. & Von Heijne, G. Genome-wide analysis of integral membrane proteins from eubacterial, archaean, and eukaryotic organisms. Protein Science 7, 1029–1038 (1998).

9 Chou, K.-C. & Elrod, D. W. Prediction of membrane protein types and subcellular locations. Proteins: Structure, Function, and Bioinformatics 34, 137–153, doi:10.1002/(sici)1097-0134(19990101)34:1<137::aid-prot11>3.0.co;2-o (1999).

10 Zhou, H. X. & Cross, T. A. Influences of membrane mimetic environments on membrane protein structures. Annu Rev Biophys 42, 361–392 (2013).

11 Seddon, A. M., Curnow, P. & Booth, P. J. Membrane proteins, lipids and detergents: not just a soap opera. Biochim Biophys Acta 3, 1–2 (2004).

12 Dowhan, W. & Bogdanov, M. Lipid-protein interactions as determinants of membrane protein structure and function. Biochemical Society transactions 39, 767–774, doi:10.1042/bst0390767 (2011).

13 Hunte, C. & Richers, S. Lipids and membrane protein structures. Current Opinion in Structural Biology 18, 406–411, doi:https://doi.org/10.1016/j.sbi.2008.03.008 (2008).

14 Palsdottir, H. & Hunte, C. Lipids in membrane protein structures. Biochimica et Biophysica Acta (BBA) - Biomembranes 1666, 2–18, doi:https://doi.org/10.1016/j.bbamem.2004.06.012 (2004).

15 Martens, C. et al. Direct protein-lipid interactions shape the conformational landscape of secondary transporters. Nat Commun 9, 018–06704 (2018).

16 Bowie, J. U. Stabilizing membrane proteins. Current Opinion in Structural Biology 11, 397–402, doi:https://doi.org/10.1016/S0959-440X(00)00223-2 (2001).

17 Duquesne, K. & Sturgis, J. N. in Heterologous Expression of Membrane Proteins: Methods and Protocols (ed Isabelle Mus-Veteau) 205–217 (Humana Press, 2010).

18 Privé, G. G. Detergents for the stabilization and crystallization of membrane proteins. Methods 41, 388–397, doi:https://doi.org/10.1016/j.ymeth.2007.01.007 (2007).

19 Shao, Z. et al. High-resolution crystal structure of the human CB1 cannabinoid receptor. Nature 540, 602, doi:10.1038/nature20613 (2016).

20 Payandeh, J., Scheuer, T., Zheng, N. & Catterall, W. A. The crystal structure of a voltage-gated sodium channel. Nature 475, 353–358, doi:10.1038/nature10238 (2011).

21 Schmidt, H. R. et al. Crystal structure of the human σ1 receptor. Nature 532, 527, doi:10.1038/nature17391 (2016).

22 Sonoda, Y. et al. Benchmarking Membrane Protein Detergent Stability for Improving Throughput of High-Resolution X-ray Structures. Structure 19, 17–25, doi:https://doi.org/10.1016/j.str.2010.12.001 (2011).

23 Phillips, R., Ursell, T., Wiggins, P. & Sens, P. Emerging roles for lipids in shaping membrane-protein function. Nature 459, 379, doi:10.1038/nature08147 (2009).

24 Andersen, O. S. & Roger E. Koeppe, I. Bilayer Thickness and Membrane Protein Function: An Energetic Perspective. Annual Review of Biophysics and Biomolecular Structure 36, 107–130, doi:10.1146/annurev.biophys.36.040306.132643 (2007).

25 Alvadia, C. et al. Cryo-EM structures and functional characterization of the murine lipid scramblase TMEM16F. eLife 8, e44365, doi:10.7554/eLife.44365 (2019).

26 Qasim, A. et al. Investigation of a KcsA Cytoplasmic pH Gate in Lipoprotein Nanodiscs. ChemBioChem 20, 813–821, doi:10.1002/cbic.201800627 (2019).

27 Pao, S. S., Paulsen, I. T. & Saier, M. H. Major Facilitator Superfamily. Microbiology and Molecular Biology Reviews 62, 1–34 (1998).

28 Reddy, V. S., Shlykov, M. A., Castillo, R., Sun, E. I. & Saier Jr, M. H. The major facilitator superfamily (MFS) revisited. The FEBS Journal 279, 2022–2035, doi:10.1111/j.1742-4658.2012.08588.x (2012).

29 Yan, N. Structural Biology of the Major Facilitator Superfamily Transporters. Annual Review of Biophysics 44, 257–283, doi:10.1146/annurev-biophys-060414-033901 (2015).

30 Marger, M. D. & Saier, M. H., Jr. A major superfamily of transmembrane facilitators that catalyse uniport, symport and antiport. Trends Biochem Sci 18, 13–20 (1993).

31 Jardetzky, O. Simple allosteric model for membrane pumps. Nature 211, 969–970 (1966).

32 Abramson, J. et al. Structure and Mechanism of the Lactose Permease of <em>Escherichia coli</em>. Science 301, 610–615, doi:10.1126/science.1088196 (2003).

33 Forrest, L. R., Kramer, R. & Ziegler, C. The structural basis of secondary active transport mechanisms. Biochim Biophys Acta 2, 167–188 (2011).

34 Quistgaard, E. M., Low, C., Guettou, F. & Nordlund, P. Understanding transport by the major facilitator superfamily (MFS): structures pave the way. Nat Rev Mol Cell Biol. 17, 123–132. doi: 110.1038/nrm.2015.1025. Epub 2016 Jan 1013. (2016).

35 Rubio-Aliaga, I. & Daniel, H. Peptide transporters and their roles in physiological processes and drug disposition. Xenobiotica 38, 1022–1042, doi:10.1080/00498250701875254 (2008).

36 Brandsch, M. Transport of drugs by proton-coupled peptide transporters: pearls and pitfalls. Expert Opinion on Drug Metabolism & Toxicology 5, 887–905, doi:10.1517/17425250903042292 (2009).

37 Giacomini, K. M. et al. Membrane transporters in drug development. Nat Rev Drug Discov 9, 215–236 (2010).

38 Dantzig, A. H. & Bergin, L. Uptake of the cephalosporin, cephalexin, by a dipeptide transport carrier in the human intestinal cell line, Caco-2. Biochim Biophys Acta 7, 211–217 (1990).

39 Biegel, A. et al. Three-dimensional quantitative structure-activity relationship analyses of beta-lactam antibiotics and tripeptides as substrates of the mammalian H+/peptide cotransporter PEPT1. J Med Chem 48, 4410–4419 (2005).

40 Daniel, H. Molecular and Integrative Physiology of Intestinal Peptide Transport. Annual Review of Physiology 66, 361–384, doi:10.1146/annurev.physiol.66.032102.144149 (2004).

41 Boggavarapu, R., Jeckelmann, J.-M., Harder, D., Ucurum, Z. & Fotiadis, D. Role of electrostatic interactions for ligand recognition and specificity of peptide transporters. BMC Biology 13, 58, doi:10.1186/s12915-015-0167-8 (2015).

42 Doki, S. et al. Structural basis for dynamic mechanism of proton-coupled symport by the peptide transporter POT. Proc Natl Acad Sci U S A 110, 11343–11348 (2013).

43 Fowler, Philip W. et al. Gating Topology of the Proton-Coupled Oligopeptide Symporters. Structure 23, 290–301, doi:https://doi.org/10.1016/j.str.2014.12.012 (2015).

44 Guettou, F. et al. Selectivity mechanism of a bacterial homolog of the human drug-peptide transporters PepT1 and PepT2. Nat Struct Mol Biol 21, 728–731 (2014).

45 Guettou, F. et al. Structural insights into substrate recognition in proton-dependent oligopeptide transporters. EMBO Rep 14, 804–810 (2013).

46 Huang, C. Y. et al. In meso in situ serial X-ray crystallography of soluble and membrane proteins at cryogenic temperatures. Acta Crystallogr D Struct Biol 72, 93–112 (2016).

47 Lyons, J. A. et al. Structural basis for polyspecificity in the POT family of proton-coupled oligopeptide transporters. EMBO Rep 15, 886–893 (2014).

48 Martinez Molledo, M., Quistgaard, E. M., Flayhan, A., Pieprzyk, J. & Low, C. Multispecific Substrate Recognition in a Proton-Dependent Oligopeptide Transporter. Structure 26, 467–476 (2018).

49 Martinez Molledo, M., Quistgaard, E. M. & Low, C. Tripeptide binding in a proton-dependent oligopeptide transporter. FEBS Lett 592, 3239–3247 (2018).

50 Minhas, G. S. et al. Structural basis of malodour precursor transport in the human axilla. eLife 3, 34995 (2018).

51 Minhas, G. S. & Newstead, S. Structural basis for prodrug recognition by the SLC15 family of proton-coupled peptide transporters. Proc Natl Acad Sci U S A 116, 804–809 (2019).

52 Newstead, S. et al. Crystal structure of a prokaryotic homologue of the mammalian oligopeptide-proton symporters, PepT1 and PepT2. Embo J 30, 417–426 (2011).

53 Parker, J. L. et al. Proton movement and coupling in the POT family of peptide transporters. Proceedings of the National Academy of Sciences 114, 13182–13187, doi:10.1073/pnas.1710727114 (2017).

54 Parker, J. L. & Newstead, S. Molecular basis of nitrate uptake by the plant nitrate transporter NRT1.1. Nature 507, 68–72 (2014).

55 Quistgaard, E. M., Martinez Molledo, M. & Löw, C. Structure determination of a major facilitator peptide transporter: Inward facing PepTSt from Streptococcus thermophilus crystallized in space group P3121. PLoS One 12, e0173126, doi:10.1371/journal.pone.0173126 (2017).

56 Solcan, N. et al. Alternating access mechanism in the POT family of oligopeptide transporters. Embo J 31, 3411–3421 (2012).

57 Ural-Blimke, Y. et al. Structure of Prototypic Peptide Transporter DtpA from E. coli in Complex with Valganciclovir Provides Insights into Drug Binding of Human PepT1. Journal of the American Chemical Society 141, 2404–2412, doi:10.1021/jacs.8b11343 (2019).

58 Zhao, Y. et al. Crystal structure of the E. coli peptide transporter YbgH. Structure 22, 1152–1160 (2014).

59 Harder, D. et al. DtpB (YhiP) and DtpA (TppB, YdgR) are prototypical proton-dependent peptide transporters of Escherichia coli. Febs J 275, 3290–3298 (2008).

60 Bippes, C. A. et al. Peptide transporter DtpA has two alternate conformations, one of which is promoted by inhibitor binding. Proc Natl Acad Sci U S A 110, 30 (2013).

61 Flayhan, A. et al. Saposin Lipid Nanoparticles: A Highly Versatile and Modular Tool for Membrane Protein Research. Structure 26, 345-355.e345, doi:https://doi.org/10.1016/j.str.2018.01.007 (2018).

62 Frauenfeld, J. et al. A saposin-lipoprotein nanoparticle system for membrane proteins. Nature methods 13, 345–351, doi:10.1038/nmeth.3801 (2016).

63 Vancraenenbroeck, R., Harel, Y. S., Zheng, W. & Hofmann, H. Polymer effects modulate binding affinities in disordered proteins. Proceedings of the National Academy of Sciences 116, 19506, doi:10.1073/pnas.1904997116 (2019).

64 Mazal, H. et al. Tunable microsecond dynamics of an allosteric switch regulate the activity of a AAA+ disaggregation machine. Nature communications 10, 1438, doi:10.1038/s41467-019-09474-6 (2019).

65 Schuler, B., Soranno, A., Hofmann, H. & Nettels, D. Single-Molecule FRET Spectroscopy and the Polymer Physics of Unfolded and Intrinsically Disordered Proteins. Annual Review of Biophysics 45, 207–231, doi:10.1146/annurev-biophys-062215-010915 (2016).

66 Grossman-Haham, I., Rosenblum, G., Namani, T. & Hofmann, H. Slow domain reconfiguration causes power-law kinetics in a two-state enzyme. Proc Natl Acad Sci U S A 115, 513–518 (2018).

67 Hellenkamp, B. et al. Precision and accuracy of single-molecule FRET measurements—a multi-laboratory benchmark study. Nature methods 15, 669–676, doi:10.1038/s41592-018-0085-0 (2018).

68 Zhu, Y., He, L., Liu, Y., Zhao, Y. & Zhang, X. C. smFRET Probing Reveals Substrate-Dependent Conformational Dynamics of E. coli Multidrug MdfA. Biophysical Journal 116, 2296–2303, doi:https://doi.org/10.1016/j.bpj.2019.04.034 (2019).

69 Daniel, H., Spanier, B., Kottra, G. & Weitz, D. From Bacteria to Man: Archaic Proton-Dependent Peptide Transporters at Work. Physiology 21, 93–102, doi:10.1152/physiol.00054.2005 (2006).

70 Brandsch, M. Drug transport via the intestinal peptide transporter PepT1. Current Opinion in Pharmacology 13, 881–887, doi:https://doi.org/10.1016/j.coph.2013.08.004 (2013).

71 Brandsch, M., Knutter, I. & Leibach, F. H. The intestinal H+/peptide symporter PEPT1: structure-affinity relationships. Eur J Pharm Sci 21, 53–60 (2004).

72 Ocheltree, S. M., Shen, H., Hu, Y., Keep, R. F. & Smith, D. E. Role and relevance of peptide transporter 2 (PEPT2) in the kidney and choroid plexus: in vivo studies with glycylsarcosine in wild-type and PEPT2 knockout mice. J Pharmacol Exp Ther 315, 240–247 (2005).

73 Weitz, D. et al. Functional and structural characterization of a prokaryotic peptide transporter with features similar to mammalian PEPT1. J Biol Chem 282, 2832–2839 (2007).

74 Prabhala, B. K. et al. Several hPepT1-transported drugs are substrates of the Escherichia coli proton-coupled oligopeptide transporter YdgR. Res Microbiol 168, 443–449 (2017).

75 Beale, J. H. et al. Crystal Structures of the Extracellular Domain from PepT1 and PepT2 Provide Novel Insights into Mammalian Peptide Transport. Structure 23, 1889–1899 (2015).

76 Kim, Y. et al. Efficient Site-Specific Labeling of Proteins via Cysteines. Bioconjugate Chemistry 19, 786–791, doi:10.1021/bc7002499 (2008).

77 Ashok, Y., Nanekar, R. & Jaakola, V.-P. Defining thermostability of membrane proteins by western blotting. Protein Engineering, Design and Selection 28, 539–542, doi:10.1093/protein/gzv049 (2015).

78 Cho, K. H., Bae, H. E., Das, M., Gellman, S. H. & Chae, P. S. Improved Glucose-Neopentyl Glycol (GNG) Amphiphiles for Membrane Protein Solubilization and Stabilization. Chemistry – An Asian Journal 9, 632–638, doi:10.1002/asia.201301303 (2014).

79 Oursel, D. et al. Lipid composition of membranes of Escherichia coli by liquid chromatography/tandem mass spectrometry using negative electrospray ionization. Rapid Communications in Mass Spectrometry 21, 1721–1728, doi:10.1002/rcm.3013 (2007).

80 Hillger, F. et al. Probing Protein–Chaperone Interactions with Single-Molecule Fluorescence Spectroscopy. Angewandte Chemie International Edition 47, 6184–6188, doi:10.1002/anie.200800298 (2008).

81 Haenni, D., Zosel, F., Reymond, L., Nettels, D. & Schuler, B. Intramolecular Distances and Dynamics from the Combined Photon Statistics of Single-Molecule FRET and Photoinduced Electron Transfer. The Journal of Physical Chemistry B 117, 13015–13028, doi:10.1021/jp402352s (2013).

82 Benke, S., Nettels, D., Hofmann, H. & Schuler, B. Quantifying kinetics from time series of single-molecule Förster resonance energy transfer efficiency histograms. Nanotechnology 28, 114002, doi:10.1088/1361-6528/aa5abd (2017).

83 Ernst, H. A. et al. Ligand binding analyses of the putative peptide transporter YjdL from E. coli display a significant selectivity towards dipeptides. Biochemical and Biophysical Research Communications 389, 112–116, doi:https://doi.org/10.1016/j.bbrc.2009.08.098 (2009).

84 Parker, J. L., Mindell, J. A. & Newstead, S. Thermodynamic evidence for a dual transport mechanism in a POT peptide transporter. eLife 2, 04273 (2014).

85 Truong-Quang, B. A. & Lenne, P. F. Membrane microdomains: from seeing to understanding. Front Plant Sci 5 (2014).

86 Low, C. et al. High-throughput analytical gel filtration screening of integral membrane proteins for structural studies. Biochim Biophys Acta 6, 9 (2013).

87 Rehan, M., Mattice, W. L. & Suter, U. W. Rotational Isomeric State Models in Macromolecular Systems. (Springer Berlin Heidelberg, 1997).

88 Chen, V. B. et al. MolProbity: all-atom structure validation for macromolecular crystallography. Acta Crystallographica Section D 66, 12–21, doi:doi:10.1107/S0907444909042073 (2010).

89 Kalinin, S. et al. A toolkit and benchmark study for FRET-restrained high-precision structural modeling. Nat Methods 9, 1218–1225 (2012).

90 Lee, N. K. et al. Accurate FRET measurements within single diffusing biomolecules using alternating-laser excitation. Biophys J 88, 2939–2953 (2005).

91 Kapanidis, A. N. et al. Alternating-laser excitation of single molecules. Acc Chem Res 38, 523–533 (2005).

92 Kapanidis, A., Majumdar, D., Heilemann, M., Nir, E. & Weiss, S. Alternating Laser Excitation for Solution-Based Single-Molecule FRET. Cold Spring Harb Protoc 2, 979–987 (2015).

93 Nettels, D. et al. Single-molecule spectroscopy of the temperature-induced collapse of unfolded proteins. Proceedings of the National Academy of Sciences 106, 20740–20745, doi:10.1073/pnas.0900622106 (2009).

94 Aznauryan, M. et al. Comprehensive structural and dynamical view of an unfolded protein from the combination of single-molecule FRET, NMR, and SAXS. Proc Natl Acad Sci U S A 113, 26 (2016).

95 Aznauryan, M., Nettels, D., Holla, A., Hofmann, H. & Schuler, B. Single-molecule spectroscopy of cold denaturation and the temperature-induced collapse of unfolded proteins. J Am Chem Soc 135, 14040–14043 (2013).

96 Dertinger, T. et al. Two-Focus Fluorescence Correlation Spectroscopy: A New Tool for Accurate and Absolute Diffusion Measurements. ChemPhysChem 8, 433–443, doi:10.1002/cphc.200600638 (2007).

97 Rusinova, E., Tretyachenko-Ladokhina, V., Vele, O. E., Senear, D. F. & Alexander Ross, J. B. Alexa and Oregon Green dyes as fluorescence anisotropy probes for measuring protein–protein and protein–nucleic acid interactions. Analytical biochemistry 308, 18–25, doi:https://doi.org/10.1016/S0003-2697(02)00325-1 (2002).

98 Likhachev. E.R. Dependence of Water Viscosity on Temperature and Pressure Zhurnal TekhnicheskoÏ Fiziki 73, 4, 135–136, https://doi.org/10.1134/1.1568496 (2003).

99 Vancraenenbroeck, R. & Hofmann, H. Occupancies in the DNA-Binding Pathways of Intrinsically Disordered Helix-Loop-Helix Leucine-Zipper Proteins. The Journal of Physical Chemistry B 122, 11460–11467, doi:10.1021/acs.jpcb.8b07351 (2018).

100 Vancraenenbroeck, R., Harel, Y. S., Zheng, W. & Hofmann, H. Polymer effects modulate binding affinities in disordered proteins. Proceedings of the National Academy of Sciences 116, 19506–19512, doi:10.1073/pnas.1904997116 (2019).

101 Schuler, B. Application of single molecule Förster resonance energy transfer to protein folding. Methods in molecular biology (Clifton, N.J.) 350, 115–138, doi:10.1385/1-59745-189-4:115 (2007).

102 Müller, B. K., Zaychikov, E., Bräuchle, C. & Lamb, D. C. Pulsed Interleaved Excitation. Biophysical Journal 89, 3508–3522, doi:https://doi.org/10.1529/biophysj.105.064766 (2005).

103 Hagen, H. Understanding disordered and unfolded proteins using single-molecule FRET and polymer theory. Methods and Applications in Fluorescence 4, 042003 (2016).

