## Supplementary information for "The conformational distribution of a major facilitator superfamily peptide transporter is modulated by the membrane composition"

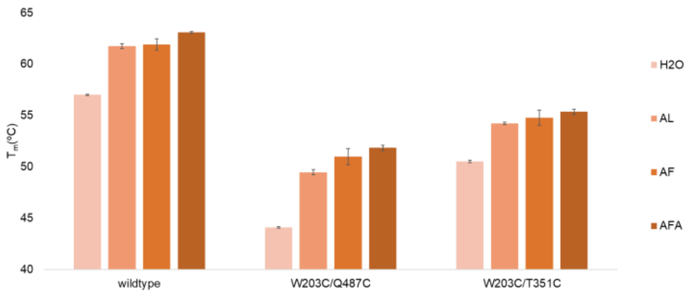


Figure S1: Thermal stability determined by nano-DSF of the DtpA variants WQ and WT, and the wildtype in the presence of the substrates Ala-Leu (AL), Ala-Phe (AF) and Ala-Phe-Ala (AFA). Error bars represent standard deviations of triplicates.

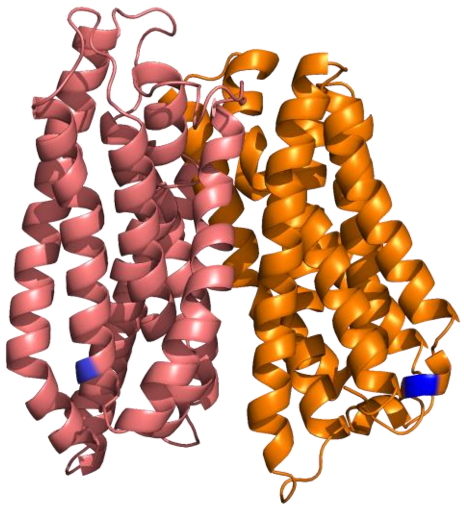

Figure S2: Labeling positions (blue) of DtpA variant WT (W203C/ T351C) on the N-terminal domain (orange) and the C- terminal domain (pink).

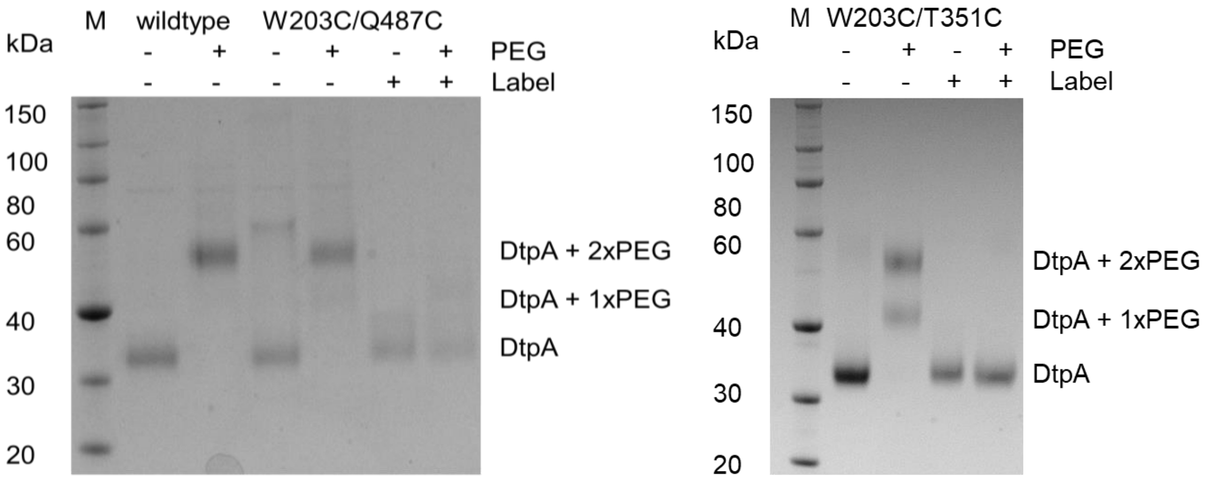


Figure S3: Cysteine accessibility test - PEGylation assay of DtpA wildtype and variants WQ and WT. Labeled ("+" with Alexa Fluor maleimide) and unlabeled ("-") proteins were incubated with PEG-maleimide to visualize the presence or absence of dye-labeling and PEG-maleimide addition, respectively. The attachment of PEG-maleimide to the free cysteine residues leads to a shift of the protein bands on SDS-PAGE to higher molecular weight (interpretation on the right). Thus, the WQ and WT cysteines were accessible for PEG-maleimide labeling via two sites, and were not accessible after labeling with the fluorescent dyes.

Table S1: $K_D$ values of N00 binding to wildtype DtpA, labeled WQ & WT, and the unlabeled WQ & WT variants, determined by the ratio of the dissociation to association rate as obtained by biolayer interferometry (Octet). Biotinylated N00 was loaded on Streptavidin biosensors and the binding was assessed at 200 nM, 100 nM, 50 nM, 25 nM, 12.5 nM and 6.25 nM DtpA and its variants. The analysis was performed using the Data Analysis software v.10.0.3.1 (fortéBIO) assuming a 1:1 stoichiometry of the protein-N00 complex.

| **Sample** | $K_D$ |
|---|---|
| Wildtype | 7.13 ± 0.03 nM |
| W203C/T351C labeled | 6.31 ± 0.05 nM |
| W203C/Q487C | 6.28 ± 0.03 nM |
| W203C/Q487C labeled | 5.40 ± 0.03 nM |

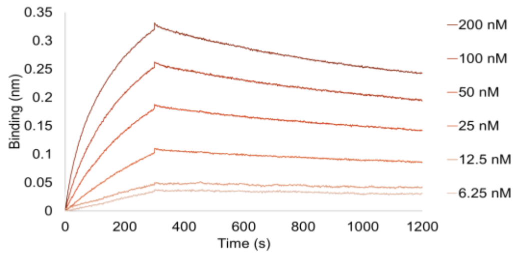


Figure S4: Binding affinity of wildtype DtpA to N00 determined by biolayer interferometry – Octet.

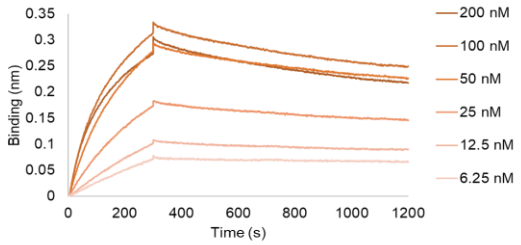


Figure S5: Binding affinity of WT variant to N00 determined by biolayer interferometry – Octet.

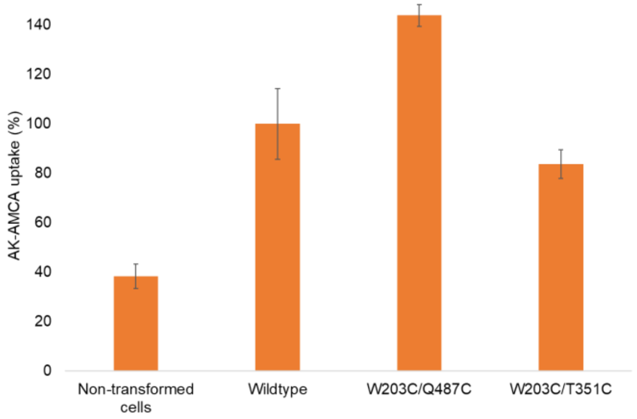


Figures S6: *In vivo* uptake assay of AK-AMCA for non-transformed cells, wildtype DtpA and the variants WQ and WT. Error bars represent the standard deviation of triplicates.

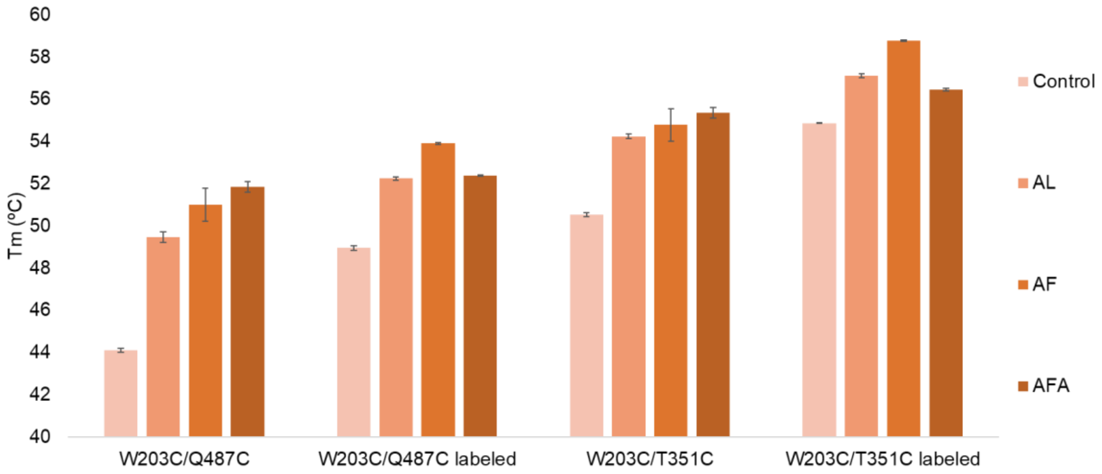


Figure S7: Thermal stability measured by nanoDSF of labeled and unlabeled variants of WQ and WT in the presence of the substrates Ala-Leu (AL), Ala-Phe (AF) and Ala-Phe-Ala (AFA). Error bars represent the standard deviation of triplicates.

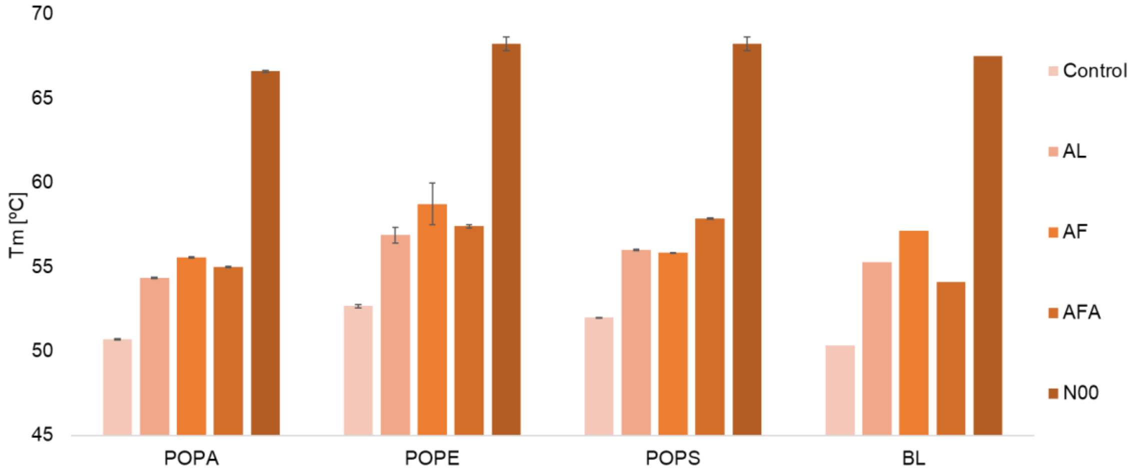


Figure S8: Thermal stability measured by nanoDSF of WQ in SapNPs of various lipid compositions: 1-palmitoyl-2-oleoyl-sn-glycero-3-phospho-L-alanine (POPA), 1-Palmitoyl-2-oleoyl-sn-glycero-3-phosphoethanol-amine (POPE), 1-palmitoyl-2-oleoyl-sn-glycero-3-phospho-L-serine (POPS) and brain-lipids extract (BL) in the presence of the the same ligands as in Fig. S7 and N00.

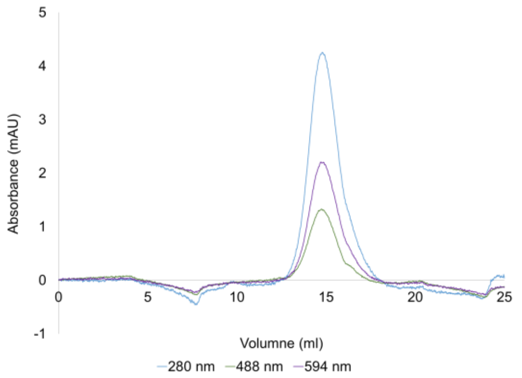


Figures S9: The elution profile (SEC) of WQ reconstituted into POPE SapNPs, run on a Superdex 200 Increase 10/300 GL column (GE Healthcare Life Sciences).

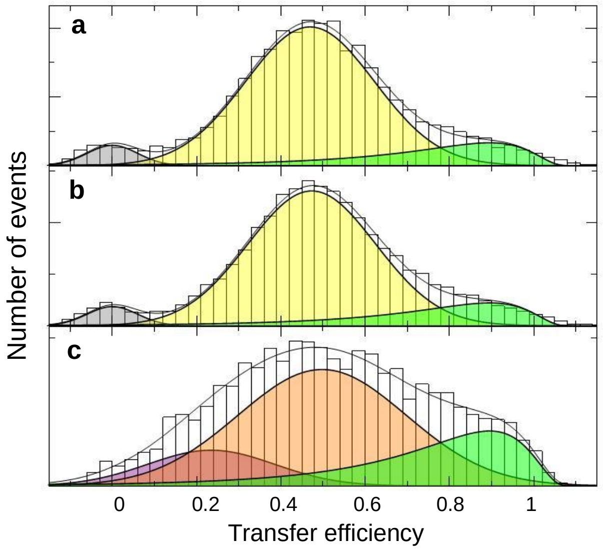


Figure S10: SmFRET histograms of the WT variant in LMNG without (a) and with (b) 8 µM N00 and WT reconstituted in POPS SapNPs (c). Solid lines are fits with a superposition of Gaussian and log-normal functions. The color code is identical to Fig. 2 and 3.

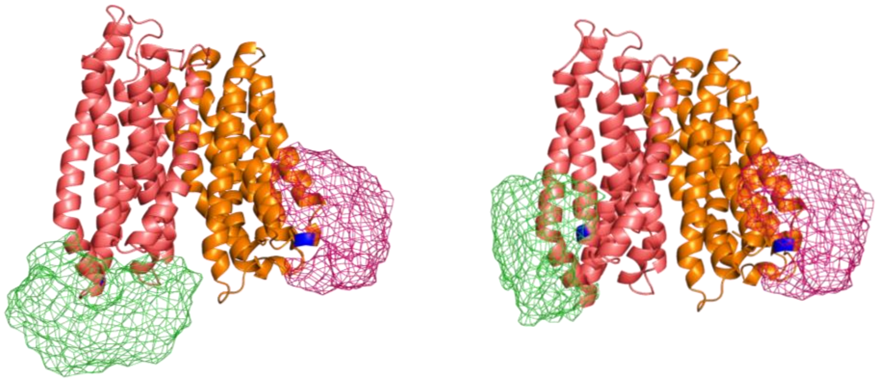

Figure S11: WQ (left) and WT (right) variants with FRET dye occupancy clouds (mesh) simulated with a Rotational Isomeric State Model. The simulation predicted mean FRET efficiency of 0.54 and 0.47 for WQ and WT respectively.

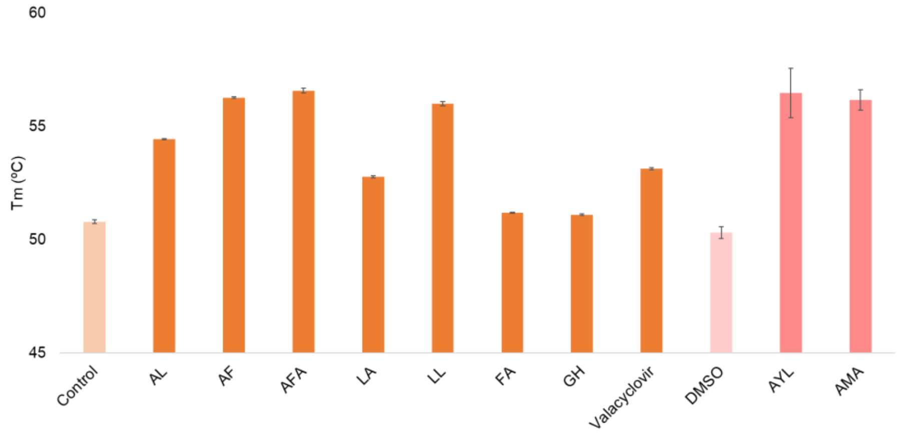


Figure S12: Thermal stability measured by nanoDSF of WQ in the presence of 2.5 mM of the following ligands: Ala-Leu (AL), Ala-Phe (AF) and Ala-Phe-Ala (AFA), Leu-Ala (LA), Leu-Leu (LL), Phe-Ala (FA), Gly-His (GH), valacyclovir, Ala-Tyr-Leu (AYL) and Ala-Met-Ala (AMA). AL, AF, AFA, LA, LL, FA, GH and valacyclovir were pre-dissolved in water, AYL and AMA in 100% DMSO.

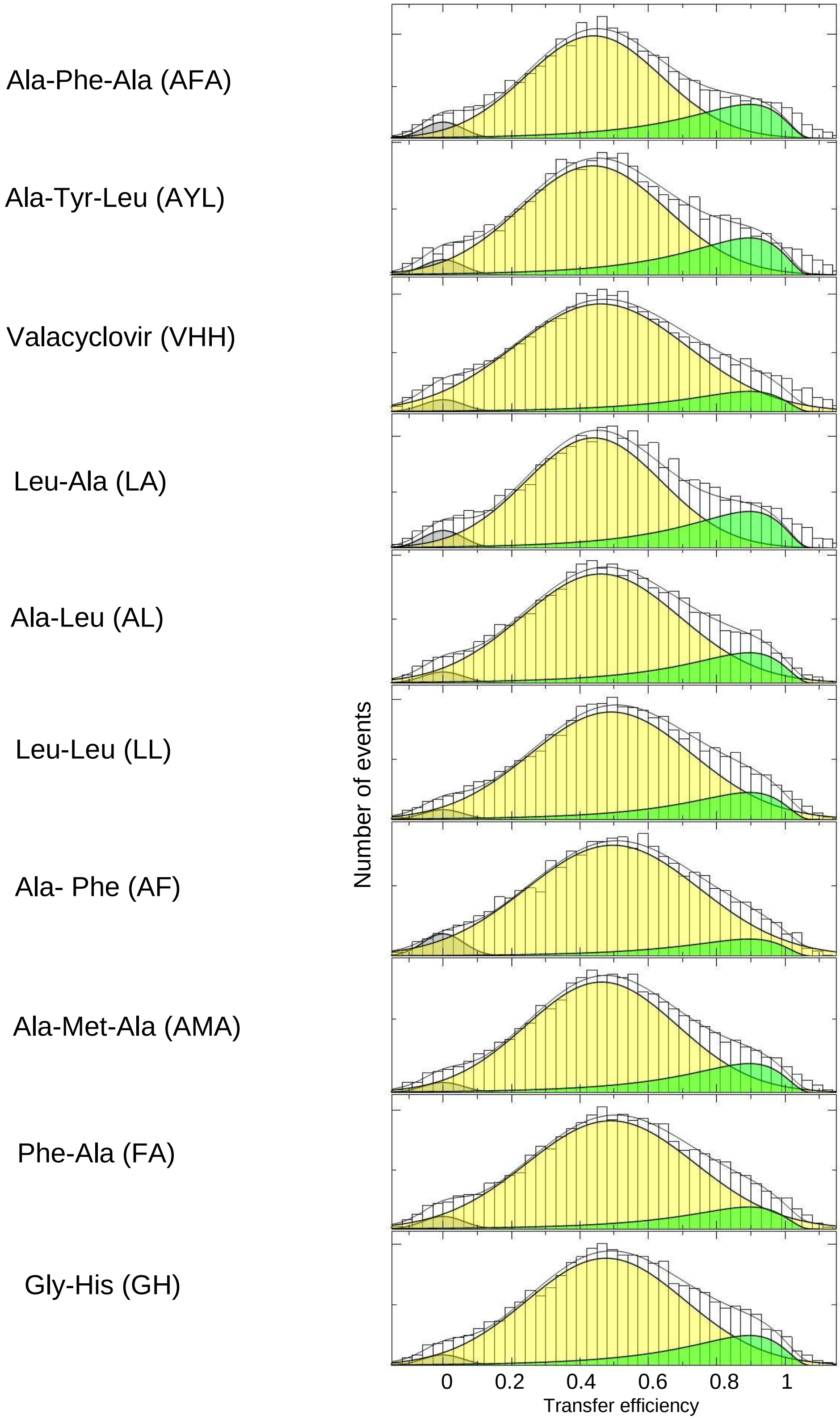


Figure S13: SmFRET histograms of WQ in LMNG with 2 mM substrates (indicated on the left). Solid lines are fits with a superposition of Gaussian and log-normal functions. The color code is identical to Fig. 2.

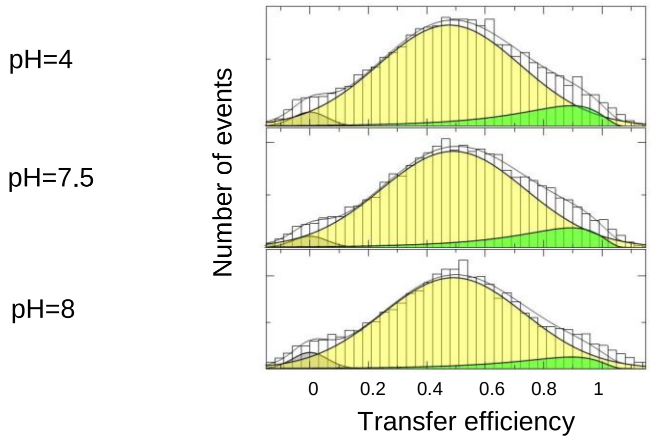


Figure S14: SmFRET histograms of WQ in LMNG at different pH values (indicated on the left). Solid lines are fits with a superposition of Gaussian and log-normal functions. The color code is identical to Fig. 2.

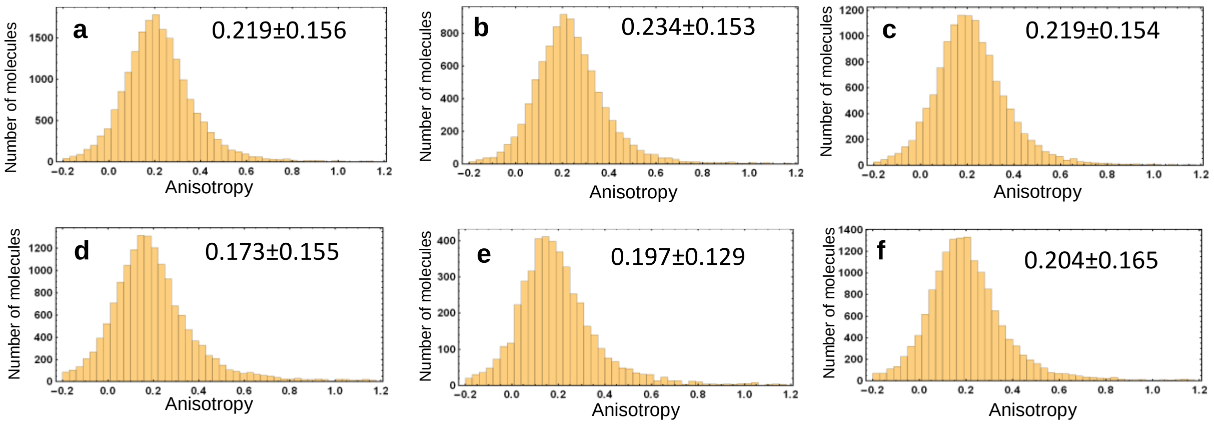


Figure S15: FRET anisotropy of the FRET labeled species (excluding molecules with inactive acceptor). (a) WQ in LMNG, (b) WT in LMNG, (c) WQ in LMNG supplied with N00, (d) WQ in POPE SapNPs, (e) WQ in POPS SapNPs, (f) WQ in POPE SapNPs supplied with N00.

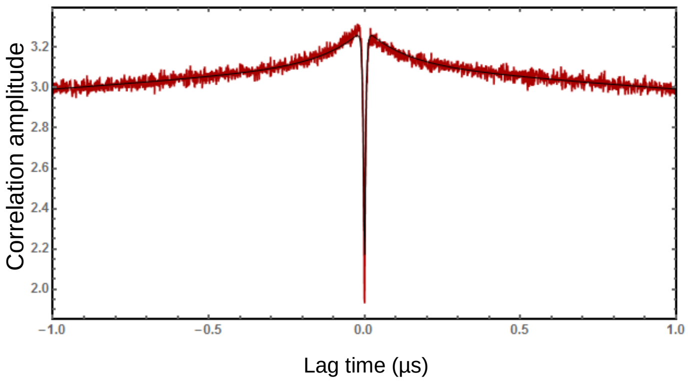


Figure S16: Nanosecond FCS (nsFCS) of the acceptor signal of WQ in POPS SapNPs after excitation at 594 nm (30uW). Solid line is a fit including components for anti-bunching (3.8 ns), the correlated decay due to quenching (127 ns, amplitude 8.1%), and triplet blinking of the dye (3.5 μs).

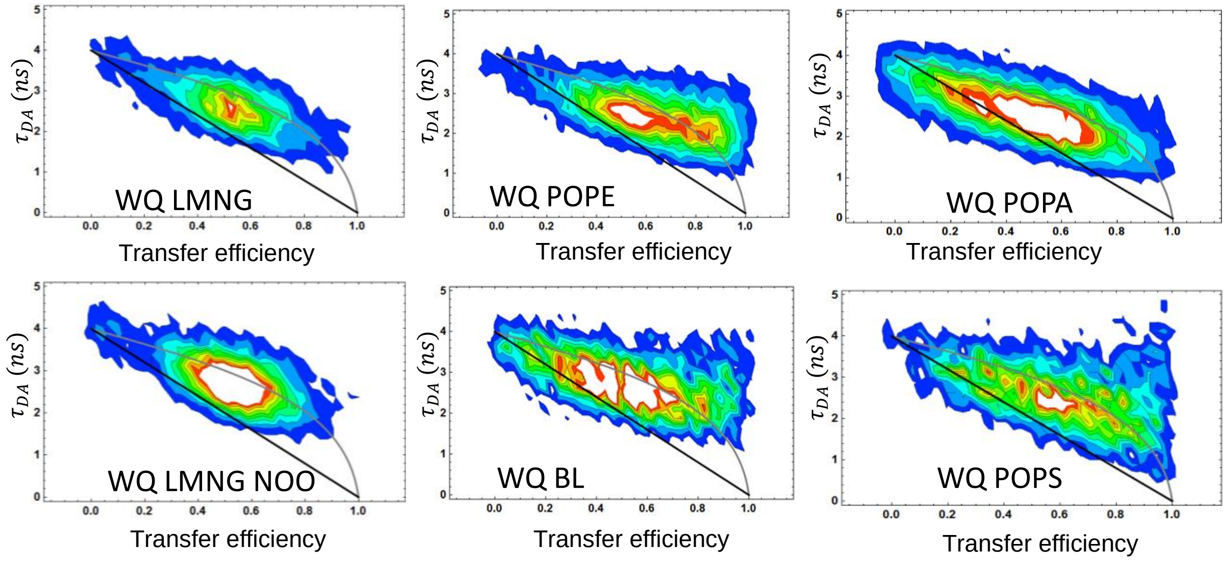


Figure S17: 2D-histograms of donor lifetime $\tau_{DA}$ vs. transfer efficiency of WQ in LMNG and reconstituted in various SapNPs (indicated). Black line is the prediction for a static distance, gray is the prediction for a Gaussian chain model.

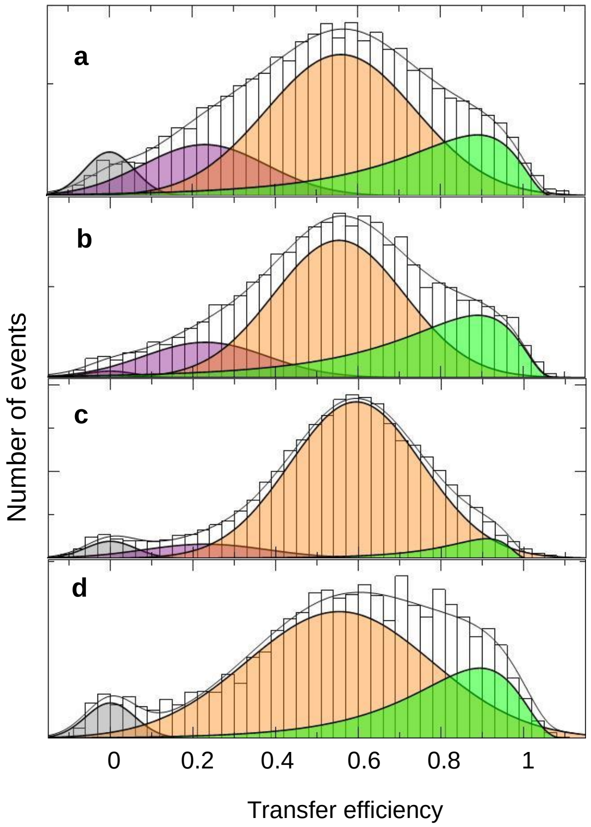


Figure S18: SmFRET histograms of WQ reconstituted in SapNPs of (a) POPS with 2mM Ala-Phe (AF), (b) POPS with 2mM Leu-Leu-Ala (LLA), (c) POPA with 2mM LLA, and (d) POPE with 2mM LLA. Solid lines are fits with a superposition of Gaussian and log-normal functions. The color code is identical to Fig. 2 and 3.

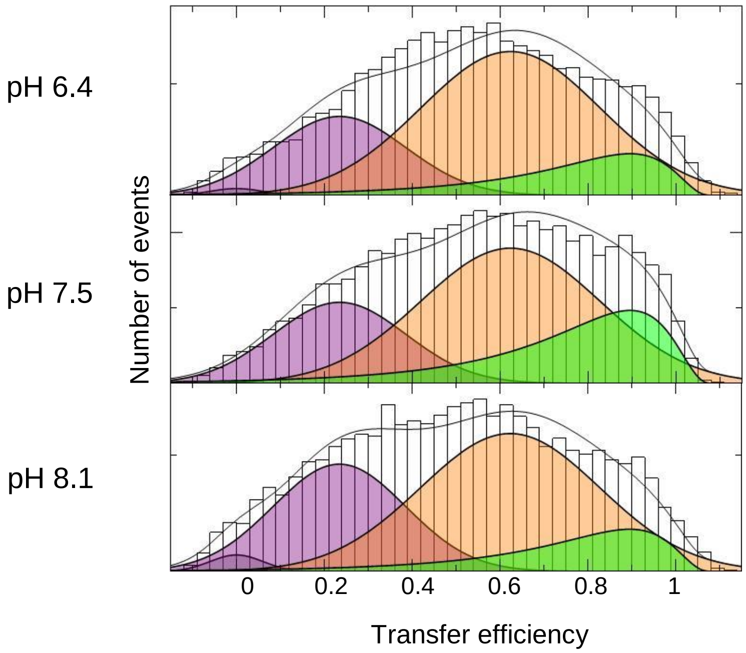


Figure S19: SmFRET histograms of WQ in POPS SapNPs at different pH values (indicated on the left). Solid lines are fits with a superposition of Gaussian and Log-normal functions. The color code is identical to Fig. 3.

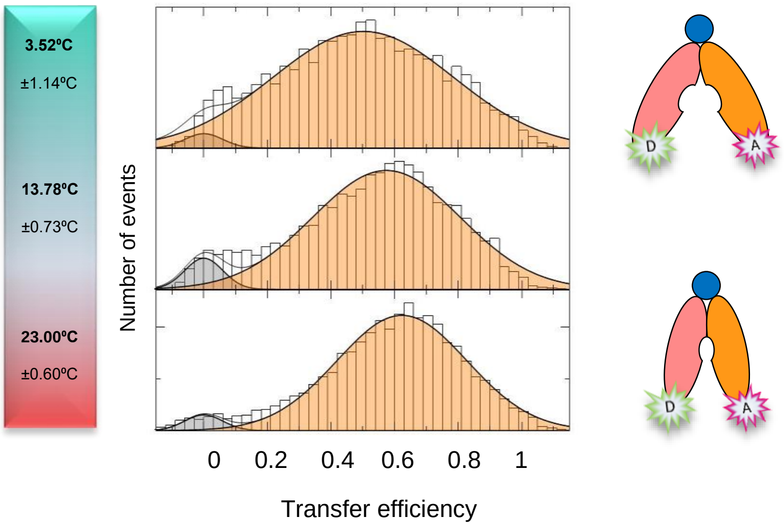


S20: SmFRET histograms of WQ reconstituted in POPS SapNPs in the presence of 8 µM nanobody N00, as a function of temperature (indicated on the left). Solid lines are fits with Gaussian peaks. Scheme (right) exemplifies the interpretation of the experiment. N00 is represented by a blue sphere. The domains are coloured as in Fig. 2a (main text).

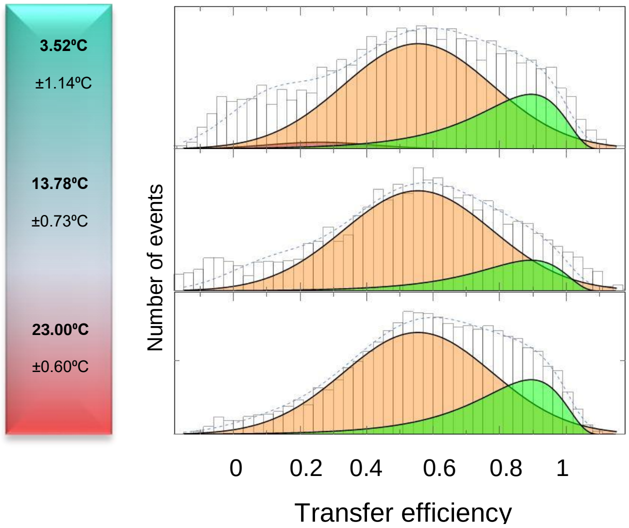


Figure S21: SmFRET histograms of the WQ variant in membrane mimicking POPE SapNPs as a function of temperature (indicate on the left). Solid lines are fits with a superposition of Gaussian and Log-normal functions. The color code is identical to Fig. 2 and 3.

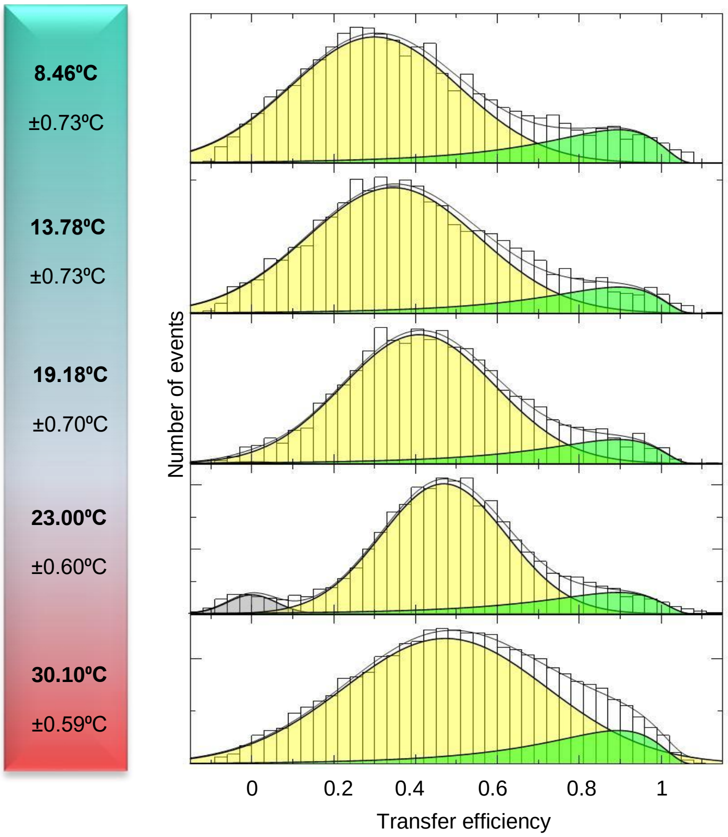


Figure S22: SmFRET histograms of WT variant in LMNG, as a function of temperature (indicated). Solid lines are fits with a superposition of Gaussian and log-normal functions. The color code is identical to Fig. 2.

**Wildtype DtpA Sequence**

ATGTCCACTGCAAACCAAAAACCAACTGAAAGCGTCAGTTTGAACGCTTTCAAACAACCGAAGGCGTTCTATCTCATCTTCTCGATTGAGTTATGGGAACGTTTGGTTATTACGGCCTACAAGGAATTATGGCTGTTTACCTGGTTAAACAACTGGGTATGTCTGAAGCGGATTCAATCACCCTTTTCTCTTCCTTTAGTGCCCTGGTTTATGGTCTGGTCGCTATCGGCGGCTGGTTAGGTGACAAGGTACTGGGTACTAAACGCGTAATTATGCTCGGCGCTATTGTGCTGGCGATTGGTTATGCTCTGGTTGCCTGGTCTGGTCACGACGCCGGTATCGTTTATATGGGTATGGCGGCTATTGCGGTCGGTAACGGCCTGTTTAAAGCTAACCCGTCTTCTCTGCTTTCTACATGCTATGAGAAAAACGACCCGCGTCTGGACGGTGCATTCACCATGTACTACATGTCCGTCAACATCGGCTCTTTCTTCTCTATGATTGCTACGCCGTGGCTGGCCGCGAAATACGGCTGGAGTGTTGCGTTTGCGTTGAGCGTTGTAGGCCTGCTGATCACTATCGTTAACTTCGCCTTCTGCCAACGCTGGGTTAAACAGTACGGTTCAAAACCAGACTTCGAGCCTATCAACTACCGTAACCTGCTGCTGACCATTATTGGTGTTGTGGCACTGATCGCTATCGCCACCTGGCTGCTGCACAATCAGGAAGTTGCGCGTATGGCGCTGGGCGTTGTTGCCTTCGGTATCGTGGTTATCTTCGGTAAAGAAGCCTTCGCGATGAAAGGTGCTGCGCGTCGTAAAATGATCGTTGCCTTCATCCTGATGCTCGAAGCCATTATCTTCTTCGTGCTGTACAGCCAGATGCCAACGTCACTGAACTTCTTTGCGATTCGTAACGTTGAGCACTCCATTCTGGGTCTGGCCGTAGAACCTGAGCAGTATCAGGCACTGAACCCGTTCTGGATCATCATCGGTAGTCCGATTCTGGCCGCTATCTATAACAAGATGGGCGATACCCTGCCGATGCCAACCAAGTTTGCAATCGGCATGGTGATGTGTTCTGGTGCGTTCCTGATTCTGCCGCTGGGTGCGAAATTCGCGTCTGACGCTGGTATCGTGTCTGTAAGCTGGCTGGTCGCAAGCTATGGCCTGCAGAGCATCGGGGAACTGATGATCTCTGGTCTGGGTCTGGCAATGGTTGCTCAACTCGTTCCGCAGCGTCTGATGGGCTTCATTATGGGTAGCTGGTTCCTGACCACTGCCGGTGCAAACCTGATTGGTGGTTATGTTGCGGGTATGATGGCTGTGCCGGATAACGTTACCGATCCGCTGATGTCACTGGAAGTCTATGGTCGCGTATTCTTGCAGATTGGTGTCGCTACTGCCGTTATTGCAGTACTGATGCTGCTGACCGCGCCGAAACTGCACCGCATGACGCAGGATGACGCTGCAGACAAAGCGGCGAAAGCAGCCGTAGCGGCAGAGAACCTCTACTTCCAATCGCACCATCATCACCACCATGATTACAAGGATGACGACGATAAGTGA

**W203C/Q487C variant Sequence**

ATGTCCACTGCAAACCAAAAACCAACTGAAAGCGTCAGTTTGAACGCTTTCAAACAACCGAAGGCGTTCTATCTCATCTTCTCGATTGAGTTATGGGAACGTTTGGTTATTACGGCCTACAAGGAATTATGGCTGTTTACCTGGTTAAACAACTGGGTATGTCTGAAGCGGATTCAATCACCCTTTTCTCTTCCTTTAGTGCCCTGGTTTATGGTCTGGTCGCTATCGGCGGCTGGTTAGGTGACAAGGTACTGGGTACTAAACGCGTAATTATGCTCGGCGCTATTGTGCTGGCGATTGGTTATGCTCTGGTTGCCTGGTCTGGTCACGACGCCGGTATCGTTTATATGGGTATGGCGGCTATTGCGGTCGGTAACGGCCTGTTTAAAGCTAACCCGTCTTCTCTGCTTTCTACATCGTATGAGAAAAACGACCCGCGTCTGGACGGTGCATTCACCATGTACTACATGTCCGTCAACATCGGCTCTTTCTTCTCTATGATTGCTACGCCGTGGCTGGCCGCGAAATACGGCTGGAGTGTTGCGTTTGCGTTGAGCGTTGTAGGCCTGCTGATCACTATCGTTAACTTCGCCTTCTCACAACGCTGCGTTAAACAGTACGGTTCAAAACCAGACTTCGAGCCTATCAACTACCGTAACCTGCTGCTGACCATTATTGGTGTTGTGGCACTGATCGCTATCGCCACCTGGCTGCTGCACAATCAGGAAGTTGCGCGTATGGCGCTGGGCGTTGTTGCCTTCGGTATCGTGGTTATCTTCGGTAAAGAAGCCTTCGCGATGAAAGGTGCTGCGCGTCGTAAAATGATCGTTGCCTTCATCCTGATGCTCGAAGCCATTATCTTCTTCGTGCTGTACAGCCAGATGCCAACGTCACTGAACTTCTTTGCGATTCGTAACGTTGAGCACTCCATTCTGGGTCTGGCCGTAGAACCTGAGCAGTATCAGGCACTGAACCCGTTCTGGATCATCATCGGTAGTCCGATTCTGGCCGCTATCTATAACAAGATGGGCGATACCCTGCCGATGCCAACCAAGTTTGCAATCGGCATGGTGATGTCATCTGGTGCGTTCCTGATTCTGCCG

CTGGGTGCGAAATTCGCGTCTGACGCTGGTATCGTGTCTGTAAGCTGGCTGGTCGCAAGCTATGGCCTGCAGAGCATCGGGGAACTGATGATCTCTGGTCTGGGTCTGGCAATGGTTGCTCAACTCGTTCCGCAGCGTCTGATGGGCTTCATTATGGGTAGCTGGTTCCTGACCACTGCCGGTGCAAACCTGATTGGTGGTTATGTTGCGGGTATGATGGCTGTGCCGGATAACGTTACCGATCCGCTGATGTCACTGGAAGTCTATGGTCGCGTATTCTTGCAGATTGGTGTCGCTACTGCCGTTATTGCAGTACTGATGCTGCTGACCGCGCCGAAACTGCACCGCATGACGTCGGATGACGCTGCAGACAAAGCGGCGAAAGCAGCCGTAGCGGCAGAGAACCTCTACTTCCAATCGCACCATCATCACCACCATGATTACAAGGATGACGACGATAAGTGA

**W203C/T351C variant Sequence**

ATGTCCACTGCAAACCAAAAACCAACTGAAAGCGTCAGTTTGAACGCTTTCAAACAACCGAAGGCGTTCTATCTCATCTTCTCGATTGAGTTATGGAACGTTTTGGTTATTACGGCCTACAAGGAATTATGGCTGTTTACCTGGTTAAACAACTGGGTATGTCTGAAGCGGATTCAATCACCCTTTTCTCTTCCTTTAGTGCCCTGGTTTATGGTCTGGTCGCTATCGGCGGCTGGTTAGGTGACAAGGTACTGGGTACTAAACGCGTAATTATGCTCGGCGCTATTGTGCTGGCGATTGGTTATGCTCTGGTTGCCTGGTCTGGTCACGACGCCGGTATCGTTTATATGGGTATGGCGGCTATTGCGGTCGGTAACGGCCTGTTTAAAGCTAACCCGTCTTCTCTGCTTTCTACATCGTATGAGAAAAACGACCCGCGTCTGGACGGTGCATTCACCATGTACTACATGTCCGTCAACATCGGCTCTTTCTTCTCTATGATTGCTACGCCGTGGCTGGCCGCGAAATACGGCTGGAGTGTTGCGTTTGCGTTGAGCGTTGTAGGCCTGCTGATCACTATCGTTAACTTCGCCTTCTCACAACGCTGCGTTAAACAGTACGGTTCAAAACCAGACTTCGAGCCTATCAACTACCGTAACCTGCTGCTGACCATTATTGGTGTTGTGGCACTGATCGCTATCGCCACCTGGCTGCTGCACAATCAGGAAGTTGCGCGTATGGCGCTGGGCGTTGTTGCCTTCGGTATCGTGGTTATCTTCGGTAAAGAAGCCTTCGCGATGAAAGGTGCTGCGCGTCGTAAAATGATCGTTGCCTTCATCCTGATGCTCGAAGCCATTATCTTCTTCGTGCTGTACAGCCAGATGCCAACGTCACTGAACTTCTTTGCGATTCGTAACGTTGAGCACTCCATTCTGGGTCTGGCCGTAGAACCTGAGCAGTATCAGGCACTGAACCCGTTCTGGATCATCATCGGTAGTCCGATTCTGGCCGCTATCTATAACAAGATGGGCGATACCCTGCCGATGCCATGTAAGTTTGCAATCGGCATGGTGATGTCATCTGGTGCGTTCCTGATTCTGCCGCTGGGTGCGAAATTCGCGTCTGACGCTGGTATCGTGTCTGTAAGCTGGCTGGTCGCAAGCTATGGCCTGCAGAGCATCGGGGAACTGATGATCTCTGGTCTGGGTCTGGCAATGGTTGCTCAACTCGTTCCGCAGCGTCTGATGGGCTTCATTATGGGTAGCTGGTTCCTGACCACTGCCGGTGCAAACCTGATTGGTGGTTATGTTGCGGGTATGATGGCTGTGCCGGATAACGTTACCGATCCGCTGATGTCACTGGAAGTCTATGGTCGCGTATTCTTGCAGATTGGTGTCGCTACTGCCGTTATTGCAGTACTGATGCTGCTGACCGCGCCGAAACTGCACCGCATGACGCAGGATGACGCTGCAGACAAAGCGGCGAAAGCAGCCGTAGCGGCAGAGAACCTCTACTTCCAATCGCACCATCATCACCACCATGATTACAAGGATGACGACGATAAGTGA

**Nanobody Sequence**

ATGGCCCAGGTGCAGCTGCAGGAGTCTGGAGGAGGATTGGTGCAGGCTGGGGGCTCTCTGAGACTCTCCTGTGCAGGCTCTGGCCGCACCTTCAGTAGTTATAACATGGGCTGGTTCCGGCAGGCTCCAGGGAAGGAGCGTGAGTTTGTAGGAGGTATTAGCTGGACTGGTCGTAGTGCCGACTATCCAGACTCCGTGAAGGGCCGATTCACTATCTCCAGAGACAACGCCAAGAACGCGGTGTATCTGCAAATGAACAGCCTGAAACCTGAAGACACGGCCGTTTATTACTGTGCAGCAAAGCAATACGGTAGTCGTGCTGACTACCCTTGGGATGACTATGACTACTGGGGCCAGGGGACCCAGGTCACCGTCTCCTCAGGGGCAGCGGAACCTGAAGCCTAG

**SaposinA Sequence**

ATGCACCATCATCATCATCATTCTTCTGGTGTAGATCTGGGTACCGAGAACCTGTACTT
CCAATCCATGGGATCCCTTCCCTGCGACATATGCAAAGACGTTGTCACCGCAGCTGGT
GATATGCTGAAGGACAATGCCACTGAGGAGGAGATCCTTGTTTACTTGGAGAAGACC
TGTGACTGGCTTCCGAAACCGAACATGTCTGCTTCATGCAAGGAGATAGTGGACTCC
TACCTCCCTGTCATCCTGGACATCATTAAAGGAGAAATGAGCCGTCCTGGGGAGGTGT
GCTCTGCTCTCAACCTCTGCGAGTCTTGA

Table S2: Primer design 0utline for point mutations in wildtype DtpA

| Mutation | Forward primer (5'-3') | Reverse primer (5'-3') |
|---|---|---|
| C140S | GATGTAGAAAGCAGAGAAGACG | GTATGAGAAAAACGACCCG |
| C200S | GAGAAGGCGAAGTTAACGATA | ACAACGCTGGGTTAAACAGTA |
| W203C | CAGCGTTGTGAGAAGG | CGTTAAACAGTACGGTTCAA |
| T351C | ATGGCATCGGCAGGGTAT | GTAAGTTTGCAATCGGCATG |
| C360S | GACATCACCATGCCGATTGC | ATCTGGTGCGTTCCTGATTCT |
| Q487C | CACGTCATGCGGTGCAG | CGATGACGCTGCAGACAAA |

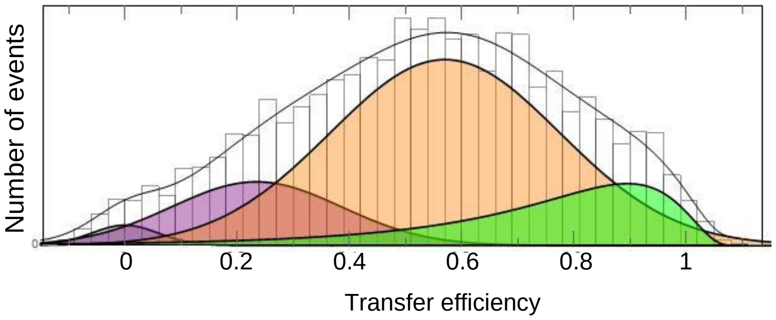


Figure S23: SmFRET histograms of the WQ variant in membrane mimicking POPS SapNPs in the absence of DTT. Solid lines are fits with a superposition of Gaussian and Log-normal functions. The color code is identical to Fig. 3.

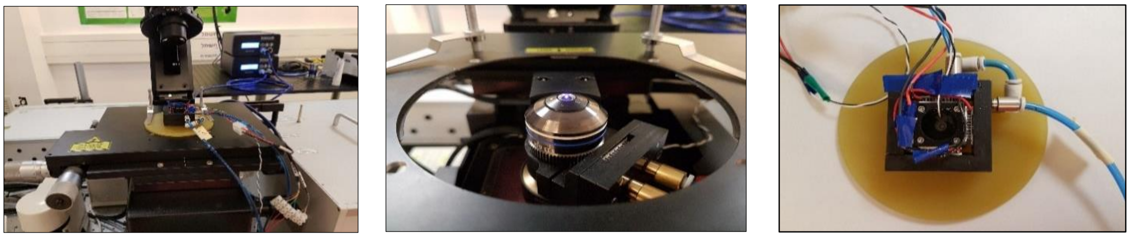

Figure S24: Temperature-controlled sample holder for single-molecule smFRET experiments. The cell adjusted over the objective (left). The objective sleeve (center), cell with a cuvette top view (right).

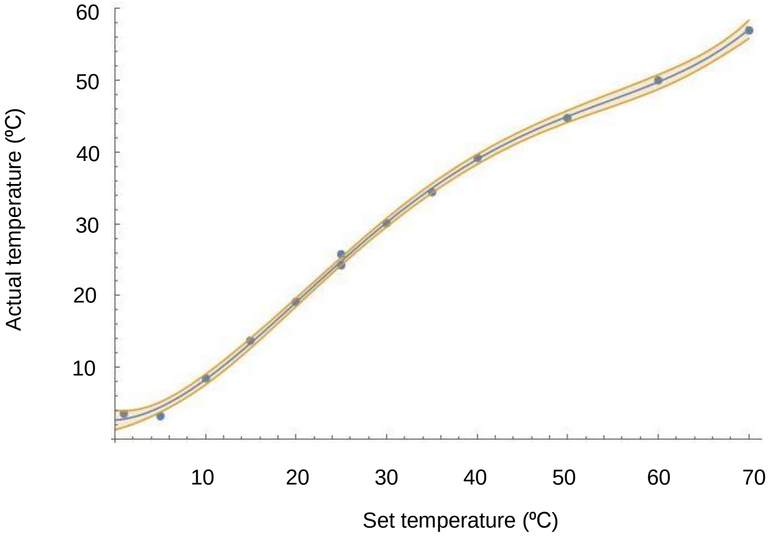


Figure S25: Calibration curve of the temperature-controlled sample holder. The calibration curve was obtained using 2f-FCS. The water viscosity was determined at different temperatures- by measuring the diffusion coefficient of Oregon Green. Viscosities were converted to temperature using the known temperature dependence of water viscosity. The solid line indicates a polynomic fit of fourth order. The collar band represents 90% confidence interval that is used for estimating the error in temperature.
